# Unbiased proteomic and forward genetic screens reveal that mechanosensitive ion channel MSL10 functions at ER-plasma membrane contact sites in *Arabidopsis thaliana*

**DOI:** 10.1101/2022.05.23.493056

**Authors:** Jennette M. Codjoe, Ryan A. Richardson, Elizabeth S. Haswell

## Abstract

Mechanosensitive (MS) ion channels are an evolutionarily conserved way for cells to sense mechanical forces and transduce them into ionic signals. The channel properties of *Arabidopsis thaliana* MscS-Like (MSL)10 have been well studied, but how MSL10 signals remains largely unknown. To uncover signaling partners of MSL10, we employed both a proteomic screen and a forward genetic screen; both unexpectedly implicated ER-plasma membrane contact sites (EPCSs) in MSL10 function. The proteomic screen revealed that MSL10 associates with multiple proteins associated with EPCSs. Of these, only VAMP-associated proteins (VAP)27-1 and VAP27-3 interacted directly with MSL10. The forward genetic screen, for suppressors of a gain-of-function *MSL10* allele (*msl10-3G, MSL10^S640L^*), identified mutations in the *synaptotagmin (SYT)5* and *SYT7* genes. We also found that EPCSs were expanded in leaves of *msl10-3G* plants compared to the wild type. Taken together, these results indicate that MSL10 can be found at EPCSs and functions there, providing a new cell-level framework for understanding MSL10 signaling. In addition, placing a mechanosensory protein at EPCS provides new insight into the function and regulation of this type of subcellular compartment.

## INTRODUCTION

Eukaryotic cells have evolved multiple mechanisms to coordinate responses between cellular compartments (Schrader et al., 2015; Mielecki et al., 2020; Sampaio et al., 2022). One such mechanism is the formation of membrane contact sites—subcellular locations where membranes of two organelles are held in close proximity by tethering proteins—which serve as sites of exchange, signaling, and organization in all eukaryotic cells (Scorrano et al., 2019; Prinz et al., 2020). One type of membrane contact site is the endoplasmic reticulum (ER)-plasma membrane (PM) contact site (EPCSs). Mammalian EPCSs are important sites for the metabolism and transport of phospholipids and allow for the coordination of ion fluxes (Zaman et al., 2020; Li et al., 2021). In plants, EPCSs help maintain phospholipid homeostasis and cell integrity (Schapire et al., 2008; Ruiz-Lopez et al., 2021), are hubs of endocytosis (Stefano et al., 2018) and autophagy (Wang et al., 2019), and regulate cell-cell transport at plasmodesmata (Levy et al., 2015; Ishikawa et al., 2020).

Several components of plant EPCSs are conserved across eukaryotes. The integral ER proteins synaptotagmins (SYTs) and vesicle-associated membrane protein (VAMP)-associated protein (VAP)27s are homologous to tricalbins and Scs2/Scs22, respectively, in yeast, and to extended-synaptotagmins and VAPs, respectively, in mammals. In yeast, tricalbins and Scs2 and Scs22 additively contribute to tethering the ER and PM to each other (Manford et al., 2012), and it is likely that plant SYTs and VAP27s also have a cooperative tethering function. Plant VAP27s may serve as a scaffold, as they are known to interact with a variety of proteins, and link EPCSs to endocytic (Stefano et al., 2018) and autophagic (Wang et al., 2019) machinery as well as to the actin cytoskeleton (Wang et al., 2014). Plant SYTs are required to maintain plasma membrane integrity in the face of stressors (Schapire et al., 2008; Yamazaki et al., 2008; Perez-Sancho et al., 2015; Ruiz-Lopez et al., 2021), probably by transporting lipids like their yeast and mammalian homologs (Saheki et al., 2016; Qian et al., 2021). Furthermore, *Arabidopsis thaliana* SYT1 changes localization and is required for cell integrity in response to mechanical pressure (Perez-Sancho et al., 2015), implicating EPCSs in the perception of mechanical stimuli. However, how mechanical information might be transmitted to or from EPCSs is completely unknown.

Organisms have evolved a variety of strategies to sense and respond to mechanical stimuli. One kind of mechanosensory protein—the mechanosensitive (MS) ion channel—represents a particularly ancient strategy that most cells still use (Árnadóttir and Chalfie, 2010; Booth et al., 2015). Most MS ion channels open and conduct ions in response to lateral membrane tension, transducing mechanical stimuli like touch, vibration, swelling, or shearing into an electrochemical signal (Kefauver et al., 2020). There is some understanding of the stimuli that activate particular plant MS channels (cell swelling, cell shrinking, encountering a barrier) as well as the adaptive processes in which they participate (relieving cell swelling, enhancing salinity tolerance, root penetration, regulating organellar morphology) (Codjoe et al., 2021). What is less understood is how signals from MS channels are coordinated across cell compartments and transduced to trigger longer-term, cell-level adaptations.

Arabidopsis MscS-Like (MSL)10 is a member of a conserved family of MS channels found in plants, bacteria, archaea, and some fungi (Hamilton et al., 2015). MSL10 is a bona fide MS ion channel and its tension-sensitive channel properties are relatively well-characterized (Haswell et al., 2008; Maksaev and Haswell, 2012; Maksaev et al., 2018). MSL10 is plasma membrane-localized (Haswell et al., 2008; Veley et al., 2014), and genetic studies have implicated it in a range of physiological roles. In response to hypo-osmotic cell swelling, MSL10 promotes a cytosolic Ca^2+^ transient, the accumulation of reactive oxygen species, the induction of *TOUCH-INDUCIBLE* gene expression, and programmed cell death (Basu and Haswell, 2020). MSL10 also contributes to systemic electrical and Ca^2+^ signaling in response to wounding (Moe-Lange et al., 2021). *MSL10* gain-of-function alleles lead to constitutive growth retardation and ectopic cell death (Basu et al., 2020) through a pathway that requires the immune co-chaperone SGT1b/RAR1/HSP90 complex, although this is likely far downstream of MSL10 activation (Basu et al., 2021). Earlier events in signal transduction by MSL10 remain unknown.

MSL10 has primarily been studied at the protein level or at the whole plant level, but its function at the subcellular level has not been addressed. To understand how MSL10 transduces mechanical information into whole plant phenotypes, we searched for potential signaling partners through forward genetic and proteomic screens. Here we describe these screens and show how both of these approaches, in combination with live-imaging assays, reveal that MSL10 functions at EPCSs.

## RESULTS

### Immunoprecipitation-mass spectrometry to identify MSL10 interactome

We first searched for signaling partners that physically interact with MSL10 using an unbiased proteomic approach. Microsomes were isolated from seedlings expressing *35S:MSL10-GFP* (Veley et al., 2014) and MSL10-GFP was immunoprecipitated from solubilized microsome extracts using GFP-Trap beads. Liquid chromatography-tandem mass spectrometry was performed on 4 replicate immunoprecipitations from *35S:MSL10-GFP* seedlings (Veley et al., 2014) as well as 4 mock immunoprecipitations from WT (Col-0) seedlings. In total, we identified 1904 peptides that mapped to 606 protein groups **(Figure 1-supplemental dataset 1).** As shown in **Figure 1a**, 239 proteins had at least 8 peptide spectral matches. Most of the proteins identified were also pulled-down with MSL10^7D^-GFP, an inactive version of MSL10 wherein 7 serines presumed to be phosphorylation sites were mutated to aspartate or glutamate (Veley et al., 2014; Basu et al., 2020 ; **Figure 1-figure supplement 1a)**. No proteins had significantly altered abundance in the MSL10 compared to MSL10^7D^ proteomes (**Figure 1-figure supplement 1b).**

**Figure 1.**
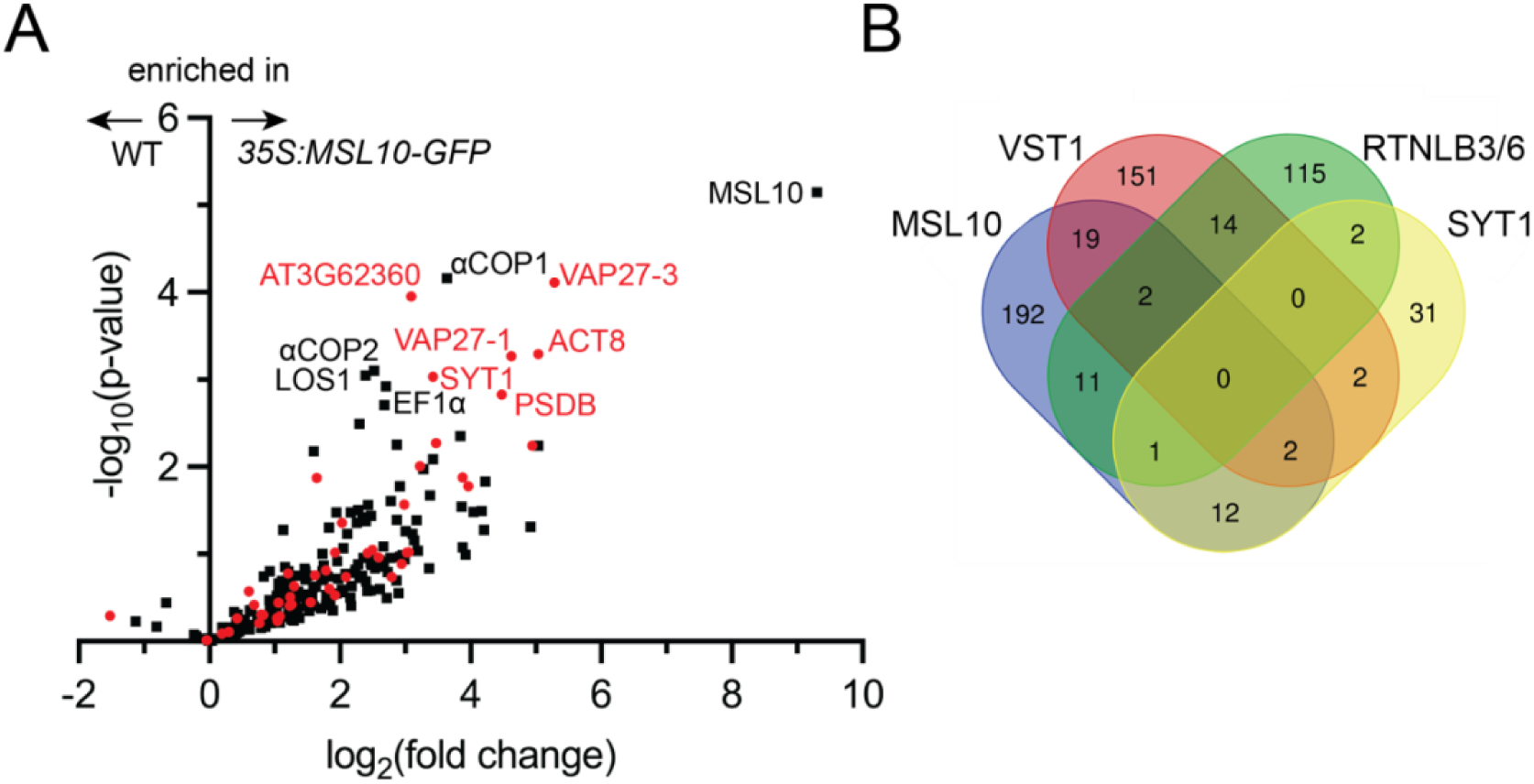
Co-immunoprecipitation-mass spectrometry identifies MSL10-GFP interactome, which shares similarities to previous EPCS interactomes. **(A)** Volcano plot showing the abundance of proteins detected in immunoprecipitations of MSL10-GFP in *35S:MSL10-GFP* seedlings (right) compared to those identified in mock immunoprecipitations using WT Col-0 seedlings (left). Proteins were identified by LC-MS/MS and the average abundance of each was quantified from the MS1 precursor ion intensities, and only those proteins with at least 8 peptide spectral matches are shown. Each protein is plotted based on its -log10(p-value) of significance based on 4 biological replicates relative to its log2(fold change) of abundance (*35S:MSL10-GFP/* WT). Proteins also detected in immunoprecipitations of EPCS proteins SYT1 (Ishikawa et al., 2020), RTNLB3/6 (Kriechbaumer et al., 2015), VST1 (Ho et al., 2016), and VAP27-1/3 (Stefano et al., 2018) are indicated in red circles; proteins unique to the MSL10 interactome are represented as black squares. The 11 most significantly enriched proteins are labelled (p-value < 0.002). (**B**) The overlap of the indicated interactomes with that of MSL10.

Among the most enriched proteins in the MSL10-GFP pull-downs were VAP27-1, VAP27-3/PVA12, and SYT1/SYTA, all of which are components of plant EPCSs (Levy et al., 2015, Wang et al., 2014, Stefano et al., 2018). This led us to perform a meta-analysis comparing the proteins that co-immunoprecipitated with MSL10 or MSL10^7D^ with four previously published interactomes of established EPCS components: SYT1 (Ishikawa et al., 2020), VAP-RELATED SUPPRESSOR OF TMM 1 (VST1) (Ho et al., 2016), reticulon-like proteins RTNLB3 and RTNLB6 (Kriechbaumer et al., 2015), and VAP27-1 and VAP27-3 (Stefano et al., 2018). 20% of the proteins that co-immunoprecipitated with MSL10-GFP have been detected at least one of these EPCS interactomes (**Figure 1a**, shown in red). And of the 10 proteins most enriched in the MSL10-GFP pull-downs (other than MSL10, the bait), 5 were previously known to be associated with plant EPCSs: SYT1, VAP27-1, VAP27-3, actin 8 (ACT8), and AT3G62360 (a predicted protein with a carbohydrate binding-like fold). Although no single protein was detected in all interactomes compared, MSL10 shared 23 interacting proteins with VST1, 15 with SYT1, and 14 with RTNLB3/6 **(Figure 1b)**. These interactomes may only partially overlap because they are incomplete, because protein complexes at EPCSs are large and difficult to fully survey, and/or because there are different EPCS complexes in different cell types or in different conditions. Nevertheless, these results indicated that MSL10 physically associates with protein complexes at EPCSs.

### MSL10 directly interacts with and co-localizes with VAP27-1 and VAP27-3

We next asked if MSL10 directly interacts with a subset of its proteome. We selected 14 of the 38 most highly enriched proteins from MSL10-GFP and/or MSL10^7D^-GFP pulldowns (fold change > 4 and p-value < 0.05), including the 5 previously associated with EPCSs for further testing. We first employed the yeast mating-based split-ubiquitin system (mbSUS) (Obrdlik et al., 2004) (**Figure 2a**). MSL10 (the bait) and the proteins being tested (the prey) were tagged with the C- and N-terminal halves of ubiquitin, respectively, such that each tag faced the cytosol. As previously reported, MSL10-Cub was able to interact with MSL10-NubG but did not interact with the potassium channel KAT1-NubG or untagged NubG (Basu et al., 2020). Of the 14 tested yeast strains, only those expressing NubG-VAP27-1 and NubG-VAP27-3 survived on minimal media when mated to yeast expressing MSL10-Cub. Consistent with our proteomic results (**Figure 1-figure 1 supplement 1b**), the interaction between MSL10 and VAP27s in the split-ubiquitin assay was not appreciably altered in MSL10 phosphovariants (**Figure 2-figure supplement 1a**), suggesting that the activation of MSL10 signaling does not alter its ability to interact with VAP27-1 and VAP27-3. Furthermore, the conserved major sperm protein domains of VAP27s were not required for interaction with MSL10 (**Figure 2-figure supplement 1b**). Along with the lack of known VAP27-binding motifs (James and Kehlenbach, 2021) in MSL10, these results indicate that MSL10 interacts with VAP27-1 and VAP27-3 in a non-canonical way.

**Figure 2.**
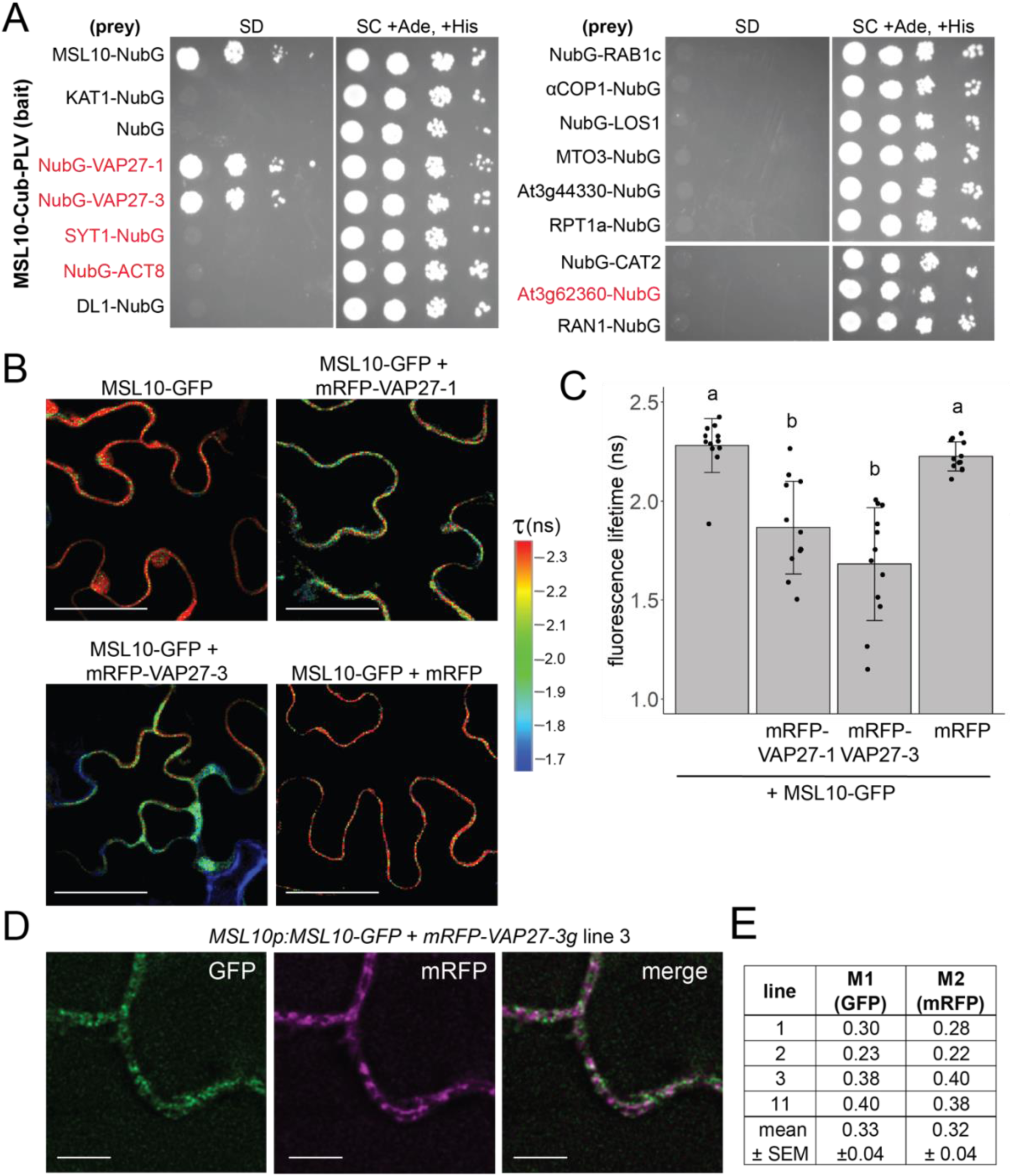
MSL10 interacts with VAP27-1 and VAP27-3. **(A)** Mating-based split-ubiquitin assay. VAMP-associated protein 27-1 (VAP27-1), VAP27-3, synaptotagmin 1 (SYT1), actin 8 (ACT8), dynamin-like (DL1), RAB GTPase homolog 1c (RAB1c), coatomer α1 subunit (αCOP1), LOW EXPRESSION OF OSMOTICALLY RESPONSIVE GENES (LOS1), METHIONINE OVERACCULATOR 3 (MTO3), AT3G44330, regulatory particle triple-A 1A (RPT1a), catalase 2 (CAT2), AT3G62360, and Ras-related nuclear protein 1 (RAN1) were tested for interaction with MSL10. Proteins labelled in red were previously detected at EPCSs. The results in (A) are consistent with another independent mbSUS assay using independent transformants. **(B-C)** *In vivo* FRET-FLIM on *UBQ:MSL10-GFP* and *UBQ:mRFP-VAP27-1* or *UBQ:mRFP-VAP27-3* transiently expressed in tobacco. **(B)** Representative heat maps of the fluorescence lifetime (τ) of GFP measured in tobacco abaxial epidermal cells 5 days post-infiltration. Scale = 50 µm. (**C**) Average GFP fluorescence lifetime. Each data point represents the value from 1 field of view (3 fields of view per plant from 4 infiltrated plants for a total of n= 12 for each combination). Error bars, SD. Groups indicated by the same letter are not statistically different according to ANOVA with Tukey’s post hoc test. (**D**) Deconvolved confocal laser scanning micrographs of leaf abaxial epidermal cells from stable Arabidopsis T1 lines co-expressing MSL10-GFP and mRFP-VAP27-3 driven by their endogenous promoters. Scale = 5 µm. **(E)** Mander’s overlap coefficients M1 and M2 calculated from images taken from 4 independent T1 lines.

We employed Fӧrster resonance energy transfer (FRET)-fluorescence lifetime imaging microscopy (FLIM) to provide additional evidence that MSL10 directly interacts with VAP27-1 and VAP27-3 in plant cells. In FRET-FLIM, when proteins are close enough for energy transfer (<10 nm), the fluorescence lifetime of the FRET donor decreases (Sun et al., 2012). MSL10-GFP transiently expressed in tobacco leaves had a fluorescence lifetime of 2.3±0.1 ns (**Figure 2b-c)**. When co-expressed with mRFP-VAP27-1 or mRFP-VAP27-3, MSL10-GFP lifetimes were 1.8±0.2 ns (a 22% decrease) and 1.6±0.3 ns (a 30% decrease), respectively. Co-expressing MSL10-GFP with free mRFP did not alter the fluorescence lifetime of GFP. These fluorescence lifetimes with and without acceptors are in the same range as those previously reported for protein-protein interactions expressed in tobacco (Wang et al., 2014, 2019).

Finally, we asked whether VAP27s and MSL10 co-localized in leaf cells of stable transgenic *Arabidopsis thaliana* lines expressing *MSL10-GFP* and *mRFP-VAP27-3* under the control of their respective promoters. We examined localization in leaf epidermal cells, where EPCSs are commonly studied and *MSL10* and *VAP27-3* are expressed (eFP Browser (Winter et al., 2007)). As expected, MSL10-GFP displayed a punctate localization at the periphery of leaf epidermal cells (**Figure 2d**) (Veley et al., 2014; Maksaev et al., 2018). In four independent *MSL10p:MSL10-GFP + mRFP-VAP27-3g* lines, mRFP signal was punctate at the cell periphery and partially co-localized with GFP signal. On average, across the four lines, 33 ± 4% of

MSL10-GFP signal co-localized with VAP27-3-mRFP, while 32 ± 4% of mRFP-VAP27-3 co-localized with MSL10-GFP (Mander’s overlap coefficient M1 and M2 respectively, **Figure 2e**).

Taken together, the data shown in Figures 1 and 2 indicate that MSL10 interacts directly with two VAP27s and indirectly with several other components of EPCSs. Because VAP27-1 and VAP27-3 are integral ER proteins (Saravanan et al., 2009; Wang et al., 2014) and MSL10 is found in the plasma membrane (Haswell et al., 2008; Veley et al., 2014), an interaction between the two would, by definition, create an EPCS.

### MSL10 alters EPCS morphology by expanding SYT1 puncta

Given that EPCS patterning is stress-responsive (Pérez-Sancho et al., 2015; Lee et al., 2019, 2020; Ruiz-Lopez et al., 2021), we hypothesized that MSL10 might serve a regulatory function at EPCSs. We began to test this hypothesis by investigating the effect of MSL10 mutant alleles on the localization of a general EPCS marker, Membrane-Attached PeriPhERal (MAPPER)-GFP (Chang et al., 2013). We crossed a *UBQ:MAPPER-GFP* line (Lee et al., 2019) to loss-of-function (*msl10-1,* (Haswell et al., 2008)) and gain-of-function (*msl10-3G,* (Zou et al., 2016; Basu et al., 2020)) *MSL10* mutant lines. In the F3 generation, we compared MAPPER-GFP localization in WT, *msl10-1* or *msl10-3G* backgrounds. MAPPER-GFP puncta looked similar in segregated WT and *msl10-1* plants (**Figure 3a-b**). In contrast, MAPPER-GFP puncta were expanded in adult *msl10-3G* plants (**Figure 3a,c**), taking up a larger proportion (13.1 ± 3.1%) of the cellular area in adult *msl10-3G* leaf epidermal cells compared to those in plants with the WT *MSL10* allele (8.7 ± 2.9%).

**Figure 3.**
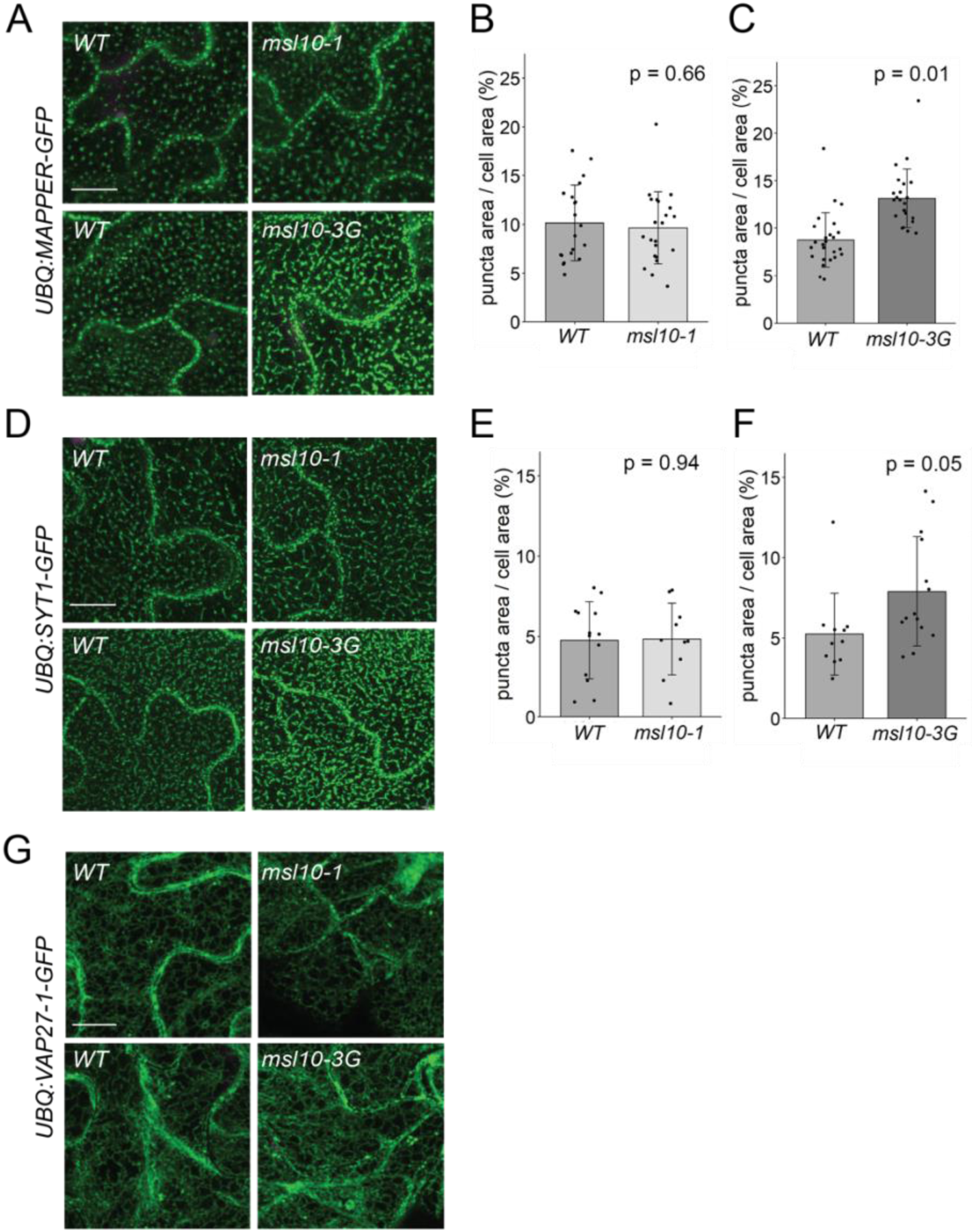
Some EPCSs are expanded in *msl10-3G* plants. Confocal maximum intensity Z-projections of GFP-tagged proteins in the indicated MSL10 backgrounds. MAPPER-GFP **(A)**, SYT1-GFP **(D)**, and VAP27-1-GFP **(G)** in 4-week-old abaxial leaf epidermal cells. Plants shown here are cousins **(A-D)** or siblings **(G)**. Green, GFP, magenta, chlorophyll autofluorescence. Scale = 10 µm. Quantification of the percentage of the leaf epidermal cell volume taken up by MAPPER-GFP **(B-C)** or SYT1-GFP **(E-F**) puncta in plants in the *msl10-1* or *msl10-3G* background compared to WT cousins. Each data point represents the mean value of 20-50 epidermal cells from one plant, n= 10-25 plants per genotype. Error bars, SD. Means were compared by Student’s t-tests when data was normally distributed **(B,E)** or Mann-Whitney U tests when it was not **(C,F)**.

We next examined VAP27 and SYT1 localization. We generated lines stably expressing VAP27-1-GFP, VAP27-3-GFP, and SYT1-GFP under control of the *UBQ10* promoter and crossed them to *msl10-1* and *msl10-3G* plants. The genotypes of surviving F2 seedlings from some of these crosses indicated genetic interactions between *MSL10* and the overexpression transgenes (**Figure 3-supplementary table 1**). For example, we were unable to isolate plants carrying the *UBQ:VAP27-3-GFP* transgene in either the *msl10-1* or *msl10-3G* backgrounds when grown on soil, and fewer *msl10-1; UBQ:SYT1-GFP* plants were isolated than would be predicted by normal Mendelian segregation (**Figure 3-supplementary table 1**).

VAP27-1-GFP is localized to the ER in Arabidopsis leaf epidermal cells, forming some puncta (although fewer than reported for VAP27-1 when transiently overexpressed in tobacco (Wang et al., 2014, 2016)). We found that the VAP27-1 localization pattern was similar in *msl10-1, msl10-3G,* and their segregated WT *MSL10* backgrounds (**Figure 3g**). As there were so few VAP27-1-GFP puncta, we did not quantify their area as for MAPPER-GFP. SYT1-GFP displayed the expected punctate localization (Levy et al., 2015; Pérez-Sancho et al., 2015). Due to the presumed synthetic lethality described above, we were unable to assess the effect of MSL10 on VAP27-3 EPCSs, and SYT1-GFP localization was unchanged in the *msl10-1* background (**Figure 3d-e**). However, in the *msl10-3G* background, SYT1-GFP puncta were expanded in leaf epidermal cells compared to the WT, leading to a modest, but significant increase in SYT1-GFP area relative to cellular area (**Figure 3d,f**). This SYT1-GFP pattern closely resembled that observed with the MAPPER-GFP marker (compare **Figure 3a** and **3d**).

### MSL10 does not contribute to EPCS rearrangement in response to osmotic perturbations

SYT-EPCSs are sensitive to environmental conditions, quickly changing localization in response to mechanical pressure (Pérez-Sancho et al., 2015) and slowly remodeling in response to freezing and salinity stress and the presence of rare ions (Lee et al., 2019, 2020; Ruiz-Lopez et al., 2021). We tested if MSL10 was required for some of these EPCS rearrangements. As previously reported (Lee et al., 2019), EPCSs marked by MAPPER-GFP in cotyledon epidermal cells expanded after a 16 hr exposure to 100 mM NaCl (**Figure 3-figure supplement 1a**). A similar MAPPER-GFP localization pattern was also observed in *msl10-1* and *msl10-3G* seedlings treated with NaCl, indicating that MSL10 does not influence the expansion of EPCSs during salinity stress. Salinity-induced EPCS expansion is reversible when seedlings are moved to media lacking NaCl, triggering a hypo-osmotic shock (Lee et al., 2019). As MSL10 plays a role in the cellular response to hypo-osmotic cell swelling (Basu and Haswell, 2020), we asked if MSL10 was also responsible for EPCS shrinking under these conditions. We found that MAPPER-GFP signal decreased in cotyledon epidermal cells 24 hr after hypo-osmotic shock (**Figure 3-figure supplement 1b**) but that this phenomenon was unaffected by the *msl10-1* or *msl10-3G* alleles. SYT1-GFP has been reported to move from a ‘beads on a string’ localization pattern to a punctate one when mechanical stress is applied (Pérez-Sancho et al., 2015). In our hands, SYT1-GFP localization always appeared punctate in cotyledon epidermal cells, and we did not see an appreciable change in this localization when pressure was added (**Figure 3-figure supplement 1c**).

### A forward genetic screen provides evidence for functional interactions between *MSL10* and *SYT5* and *SYT7*

Above, we describe physical interactions between MSL10 and the EPCS components VAP27-1 and VAP27-3, and a functional interaction wherein SYT1 EPCSs are expanded in *msl10-3G* plants. Further evidence for functional interactions between MSL10 and EPCS components came from a genetic screen that was performed at the same time as the above experiments. We used the obvious growth defect of *msl10-3G* plants (Zou et al., 2016; Basu et al., 2020) as the basis of a visual screen, as illustrated in **Figure 4a**. EMS-induced suppressor mutants, referred to as *<u>s</u>uppressed <u>d</u>eath from <u>m</u>sl10-3G* (*sdm*), were initially isolated based on increased height compared to parental *msl10-3G* plants in the M1 and M2 generations. As *msl10-3G* plants share some of the characteristics of lesion-mimicking-mutants (Basu et al., 2021), and intragenic mutations are particularly common in suppressor screens of lesion-mimicking mutants (van Wersch et al., 2016), we sequenced *MSL10* exons in all 40 mutant lines. Indeed, 35 had a missense mutation in the *MSL10* coding or splice-junction sequences (**Figure 4-figure supplement 1a**). The five remaining *sdm* mutants were presumed to have extragenic suppressor mutations. The mapping-by-sequencing strategy we employed (see below) successfully identified extragenic suppressor mutations for two of these five, *sdm26* and *sdm34*.

**Figure 4.**
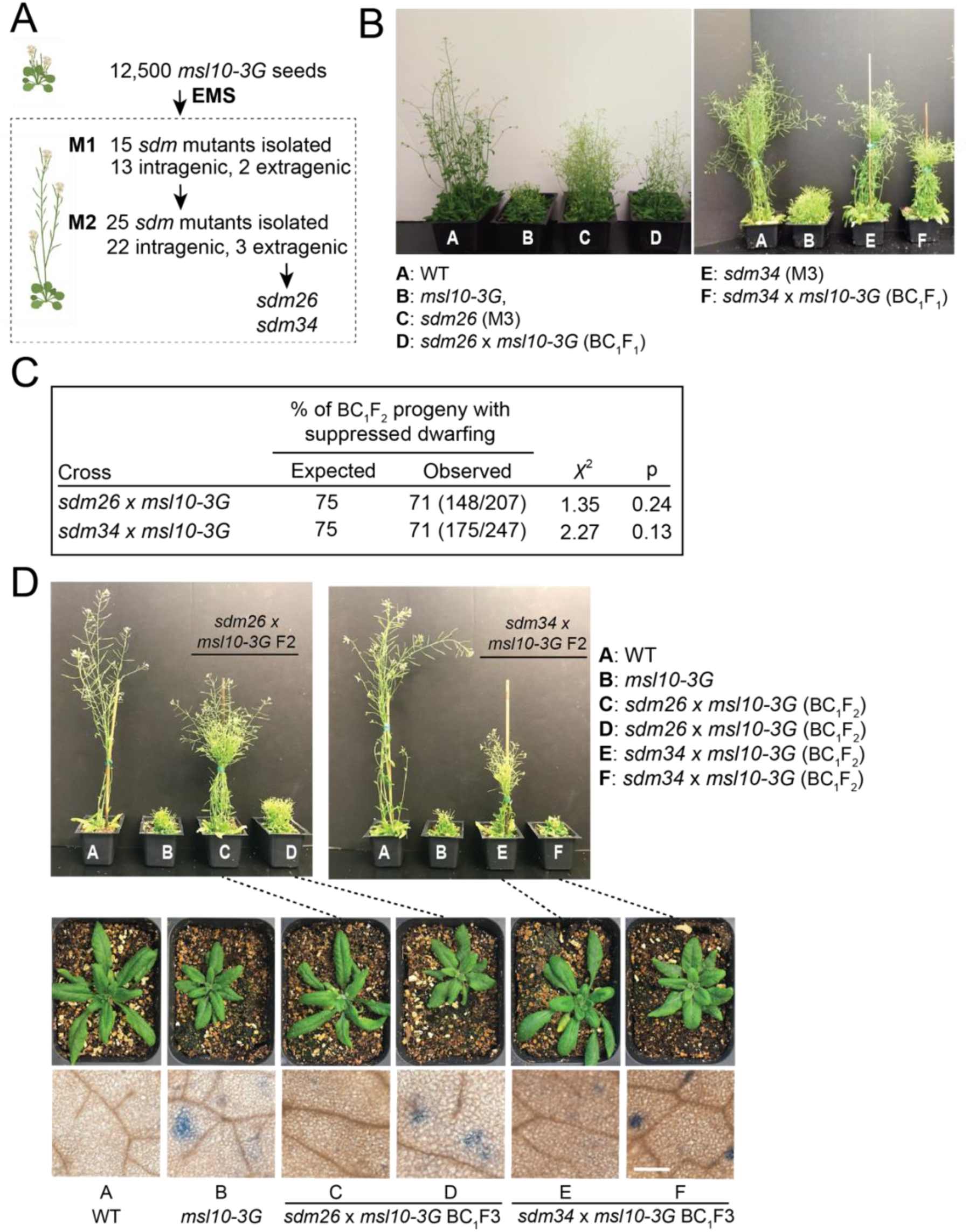
A forward genetic screen identifies *sdm26* and *sdm34,* dominant suppressors of *msl10-3G* height and ectopic cell death phenotypes, *sdm26* and *sdm34*. **(A)** Schematic of the screen. **(B)** Images of the indicated plants after 4-5 weeks of growth. **(C)** Segregation of height phenotypes in the BC_1_F2 generation, compared to the expected segregation ratio assuming the *sdm* alleles were dominant. **(D)** Siblings of backcrossed *sdm26* and *sdm34* mutants were isolated that were fixed for the *sdm* (suppressed dwarfing) or *msl10-3G* (dwarf) phenotypes. Top: 5-week-old BC_1_F_2_ plants of the indicated genotypes. Middle: 4-week-old BC_1_F_3_ progeny of plants at the top, as indicated with dashed lines. Bottom: Leaves of 4-week-old BC1F3 plants stained with Trypan blue to assess cell death. These results are representative of at least five other plants for each genotype. Scale = 300 µm.

Notably, *sdm26* and *sdm34* mutant plants were taller than *msl10-3G* plants but not as tall as WT plants (**Figure 4b**). The offspring of both *sdm26* and *sdm34* backcrosses to *msl10-3G* (BC_1_F1 plants) were as tall as their *sdm* parents (**Figure 4b**). Furthermore, in the BC_1_F2 generation, plants with intermediate height (*sdm* phenotype) were present approximately 3:1 relative to those with the *msl10-3G* dwarf phenotype (**Figure 4c**), indicating that the *sdm* mutations are dominant in the *msl10-3G* background, at least for this phenotype. When *sdm26* and *sdm34* plants were outcrossed to the *msl10-1* null allele, plants with the parental *msl10-3G* phenotype were recovered in the F2 generation (**Figure 4-figure supplement 1b**), confirming that the *sdm26* and *sdm34* lesions are extragenic alleles unlinked to *MSL10*. Another characteristic phenotype of *msl10-3G* plants, ectopic cell death, was also suppressed in *sdm26* and *sdm34* leaves compared to those of parental and segregating *msl10-3G* siblings, although the *sdm* mutants exhibited slightly more cell death than WT plants (**Figure 4d**).

We employed a whole genome sequencing strategy to identify the mutations responsible for *sdm26* and *sdm34* phenotypes (**Figure 5a**). BC_1_F2 plants were separated by phenotype into pools of 50 plants each, and genomic DNA was extracted from pooled tissue and sequenced at 80x coverage. As *sdm26* and *sdm34* are dominant suppressor mutations, we searched for EMS-induced SNPs that 1) had an allele frequency of 0.66 in the pool of plants with the *sdm* phenotype and 2) were absent in the *msl10-3G* phenotype pool. Intervals of adjacent SNPs with such allele frequencies were found on chromosome 1 for *sdm26* and chromosome 3 for *sdm34* (**Figure 5-figure supplement 1**). We failed to identify clear intervals of linked SNPs with the expected allele frequencies for the other 3 presumed extragenic mutants.

**Figure 5.**
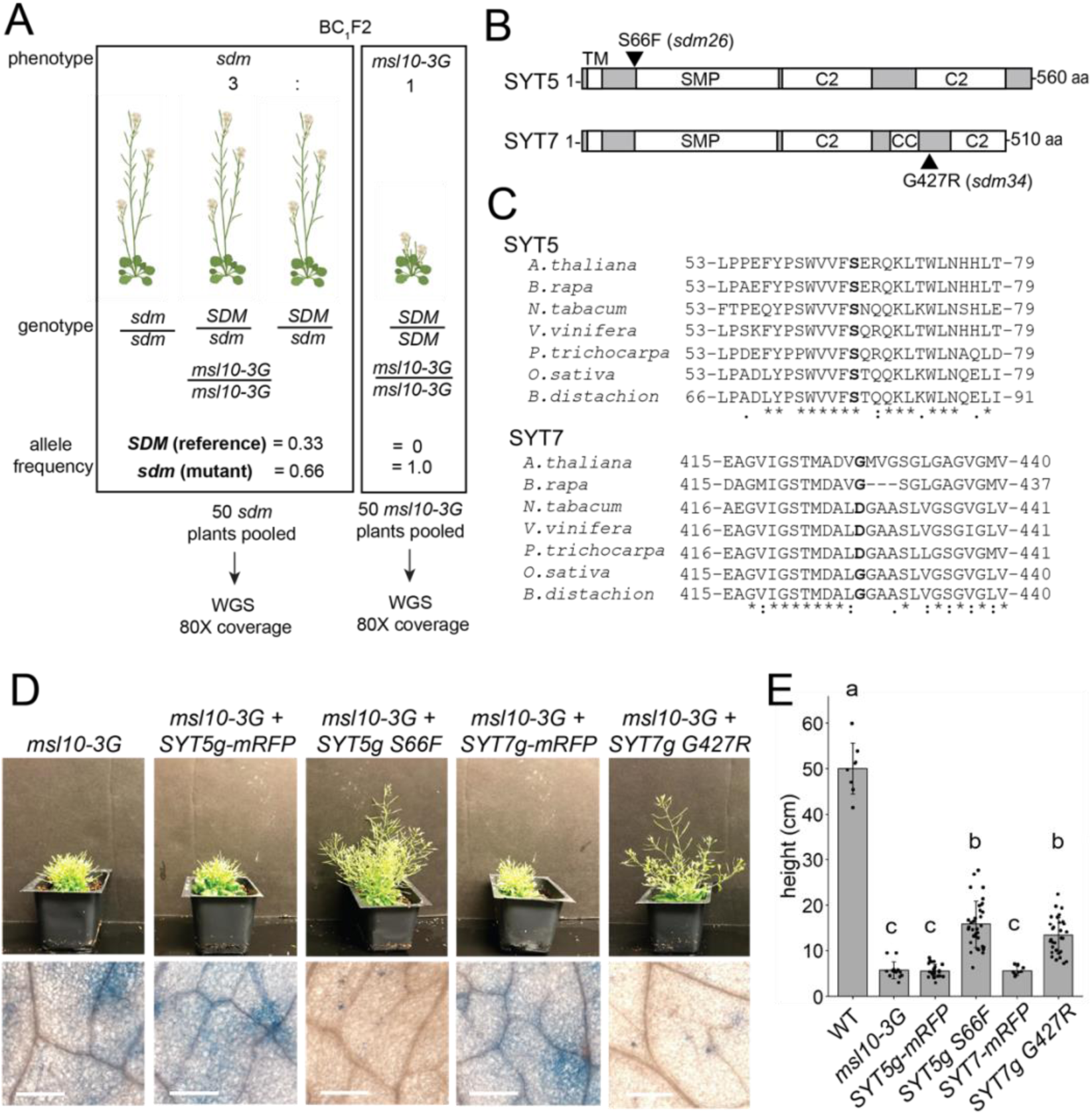
*SYT5 S66F* and *SYT7 G427R* are the causal mutations in *sdm26* and *sdm34*, respectively. **(A)** Overview of backcrossing and mapping-by-sequencing of *sdm* mutants. **(B)** Location of *sdm26* and *sdm34* missense mutations in the SYT5 and SYT7 proteins, respectively. UniProt was used to predict protein domains and their location. TM, transmembrane; SMP, synaptogamin-like mitochondrial-lipid-binding protein domain; CC, coiled coil; C2, Ca^2+^ binding. **(C)** Conservation of Ser66 and Gly427 residues in SYT5 and SYT7 homologs, respectively, in the predicted proteomes of selected seed plants. **(D-E)** Phenotypes of *msl10-3G* plants expressing WT or *sdm* mutant SYT5 and SYT7 transgenes. **(D)** Top: Images of representative T1 lines. Bottom: Trypan blue staining of a leaf from the same plants. Scale = 300 µm. **(E)** Mean and standard deviation of plant height of n= 9-32 T1 lines per construct.

The intervals in *sdm26* and *sdm34* contained 8 and 13 genes, respectively. The *sdm26* genome encoded a missense mutation (Ser66→Phe) in the *synaptotagmin 5* (*SYT5)* gene and the *sdm34* genome encoded a Gly427→Arg substitution in *synaptotagmin 7 (SYT7, CBL1, NTMC2T4;* **Figure 5b**). SYT5 and SYT7 are known to interact with each other and with SYT1 at EPCSs (Ishikawa et al., 2020; Lee et al., 2020). Given these results, and that MSL10 interacts with EPCS proteins **(Figure 1, 2)**, the SNPs in *SYT5* and *SYT7* were promising candidates for causing the suppression of the *msl10-3G* phenotypes in *sdm26* and *sdm34*. However, it remained possible that lesions elsewhere in these intervals were instead responsible.

We therefore attempted to recreate the *sdm* phenotypes by expressing *SYT5 S66F* and *SYT7 G427R* from transgenes in unmutagenized *msl10-3G* plants. We expected to see *sdm*-like phenotypes in the T1 generation because the suppressor mutations in *sdm26* and *sdm34* plants were dominant. As anticipated, *msl10-3G*+*SYT5g S66F* and *msl10-3*+*SYT7g G427R* T1 plants were taller than untransformed *msl10-3G* plants (**Figure 5d**). The amount of ectopic cell death was also suppressed compared to *msl10-3G* leaves. WT *SYT5g-mRFP* or WT *SYT7g-mRFP* had no discernable effect on plant height or ectopic cell death in T1 plants in the *msl10-3G* background. These results provide strong evidence that *SYT5 S66F* and *SYT7 G427R* mutations caused suppression of *msl10-3G* phenotypes in the *sdm26* and *sdm34* mutants, respectively.

To address whether the *sdm26* and *sdm34* mutations might be dominant negative, we crossed *msl10-3G* plants to null *syt5* and *syt7* alleles (Ishikawa et al., 2020). Double *syt5; msl10-3G* and *syt7; msl10-3G* mutants resembled *msl10-3G* plants (**Figure 6-supplemental figure 1a-b)**. The inability of null *syt5* and *syt7* alleles to suppress *msl10-3G* phenotypes indicates that the *sdm26 (SYT5 S66F)* and *sdm34 (SYT7 G427R)* alleles do not cause suppression by impairing the function of WT SYT5 or SYT7. Additionally, the null *syt1-2* allele (Ishikawa et al., 2020) had no effect on *msl10-3G* growth defects or ectopic death (**Figure 6-supplemental figure 1a**).

### *sdm26* and *sdm34* alleles do not alter SYT5 or SYT7 localization or MSL10 levels

The SYT5 S66F and SYT7 G427R point mutations occur in very different parts of the synaptotagmin proteins and are not located in any of the predicted functional domains (Ishikawa et al., 2020; Lee et al., 2020; The UniProt Consortium, 2021) (**Figure 5b**). However, S66 is fully conserved in SYT5 homologs from monocots and dicots and G427 is partially conserved in SYT7 homologs from Brassicacae and monocots (**Figure 5c**), and thus may be important for structure or function. We first investigated if the *sdm* point mutations change the localization of SYT5 and SYT7. When transiently expressed in tobacco, SYT5 S66F-mRFP and SYT7 G427R-mRFP had similar localization and dynamics to their WT counterparts, localizing to dynamic ER tubules and to puncta that persisted over time, as previously reported (Ishikawa et al., 2020; Lee et al., 2020) (**Figure 6-supplemental figure 1c; Movies 1-4**). Additionally, the *sdm* point mutations do not alter *SYT5* or *SYT7* transcript stability (**Figure 6-supplemental figure 1d**). To rule out a trivial explanation for the suppression of *msl10-3G* phenotypes—that the *sdm26* and *sdm34* alleles decrease MSL10 expression and/or stability—we examined expression of *MSL10p:MSL10-GFP* expression in those backgrounds. We found equivalent MSL10-GFP fluorescence and protein levels in *sdm26* plants and their WT siblings, and in *sdm34* plants compared to their WT siblings (**Figure 6-supplemental figure 1e-f**). In summary, the *sdm26* and *sdm34* alleles do not affect MSL10 expression or protein stability, nor SYT5 or SYT7 localization, suggesting that they suppress MSL10 signaling in some other way.

### EPCS expansion is not suppressed in *sdm26* and *sdm34* mutants

Given that SYT1-EPCSs were expanded in *msl10-3G* mutants, we wondered if increased connections between the ER and PM in *msl10-3G* plants might be responsible for the growth retardation and ectopic cell death associated with this allele. If this were the case, the enhanced EPCS area observed in *msl10-3G* plants would be suppressed by *sdm26* or *sdm34* alleles. To test this idea, we crossed *UBQ:MAPPER-GFP* plants to the *sdm26* mutant. To our surprise, the larger EPCS area in *msl10-3G* plants (13.7±4.2%) was not suppressed in *sdm26* leaf epidermal cells (13.5±3.7%) (**Figure 6a-b**). The same observation was made in plants derived from a *UBQ:MAPPER-GFP x sdm34* cross (**Figure 6c-d**). Thus, differences in ER-PM connectivity, at least as marked by MAPPER-GFP, do not drive the phenotypic differences we observe between WT, *msl10-3G,* and *sdm* plants.

**Figure 6.**
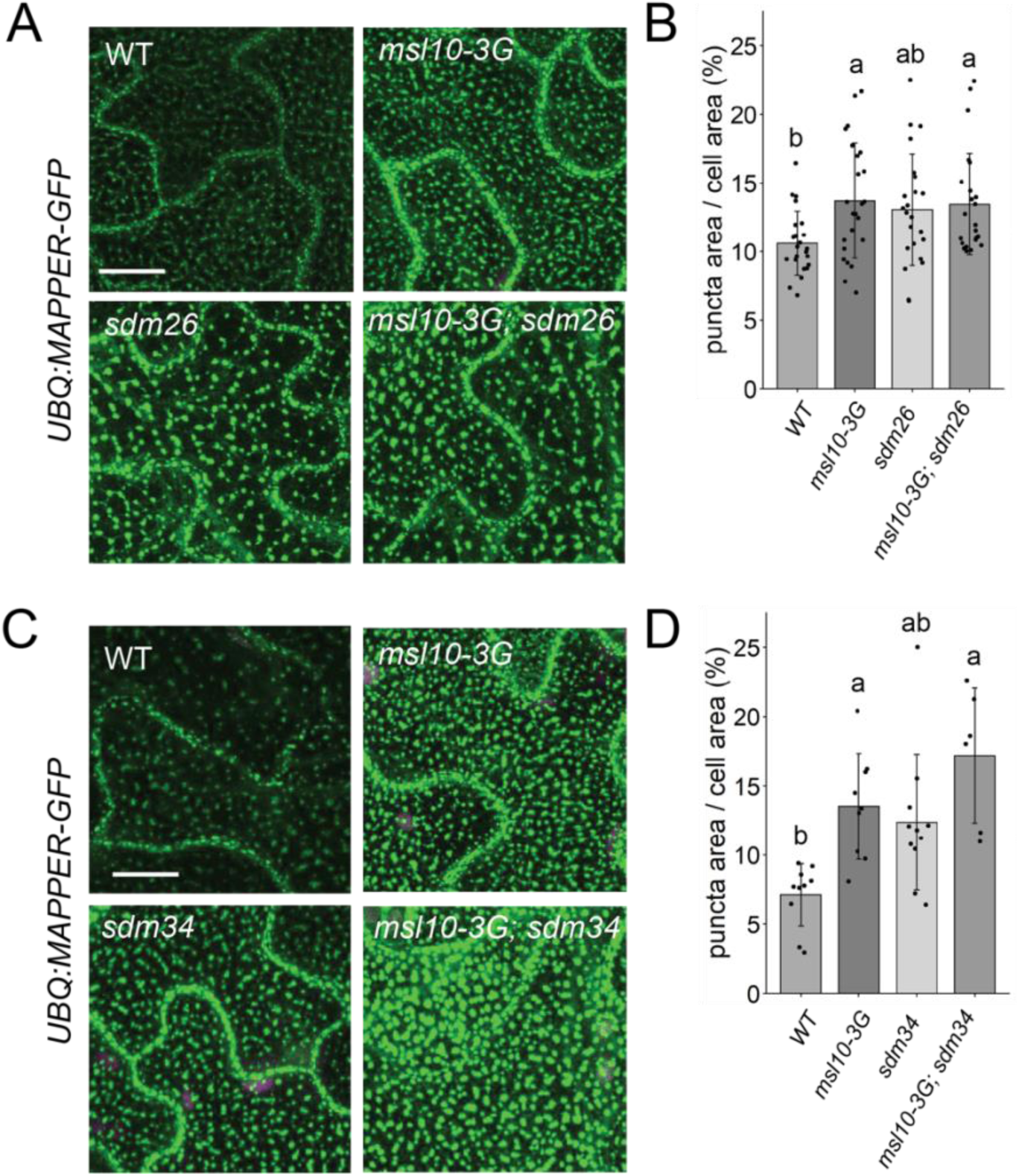
*sdm26* and *sdm34* alleles do not suppress expanded EPCSs in *msl10-3G* leaves. **(A,C)** Confocal maximum intensity Z-projections of MAPPER-GFP fluorescence in 4-week-old abaxial leaf epidermal cells of the indicated genotypes. Scale= 10 µm. **(B,D)** Quantification of the percentage of the leaf epidermal cell volume taken up by MAPPER-GFP puncta in plants of the indicated genotypes. Each data point represents the mean value of 20-50 epidermal cells from one plant, n = 6-23 plants per genotype. Error bars, SD. Groups indicated with the same letters are not significantly different as assessed by Kruskal-Wallis with Dunn’s post hoc test when measurements were not normally distributed (**B**) or ANOVA with Scheffe’s post hoc test when they were **(D)**.

### MSL10 does not interact with SYT5 or SYT7 or influence their localization

As SYT1-EPCSs were expanded in *msl10-3G* leaf epidermal cells (**Figure 3d,f**), and SYT1 can interact with SYT5 and SYT7 (Ishikawa et al., 2020; Lee et al., 2020), we asked if SYT5 and SYT7 localization were also altered in the *msl10-3G* background. We transformed WT Col-0 plants with GFP-tagged constructs under the control of the *UBQ10* promoter and crossed these lines to *msl10-1* and *msl10-3G* plants. Both SYT5-GFP and SYT7-GFP had a partially punctate, partially ER localization, as we had observed with mRFP-tagged versions expressed transiently in tobacco **(Figure 7a-b, Figure 6-supplemental figure 1**), and this localization pattern was unaffected by the *msl10-3G* or *msl10-1* alleles.

**Figure 7.**
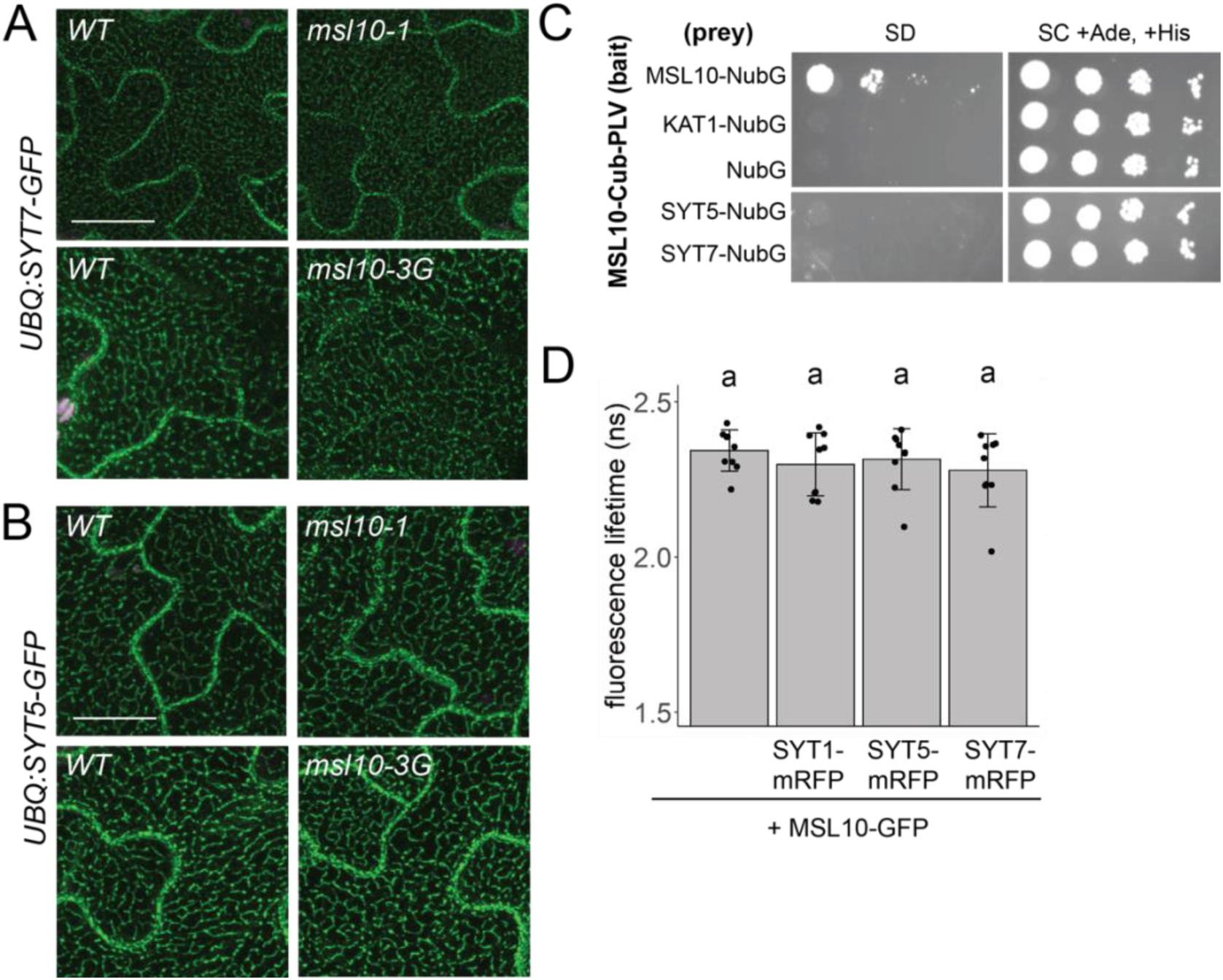
MSL10 does not interact with SYT5 or SYT7 nor alter their localization. **(A-B)** Confocal maximum Z-intensity projections of abaxial leaf epidermal cells of 4-week-old plants with the indicated *MSL10* alleles. Plants in **(A)** are F2 siblings, in **(B)** F3 cousins. Scale = 15 µm. **(C)** Mating-based split-ubiquitin assay testing the interaction of MSL10 with SYT5 and SYT7, performed as in Figure 2a. **(D)** Fluorescence lifetime (τ) of GFP measured using FRET-FLIM when *UBQ:MSL10-GFP* was transiently expressed tobacco leaves for 5 days, with or without *UBQ:SYT-mRFP* constructs. Each data point represents the value from 1 field of view (3 fields of view per plant from 3 infiltrated plants for a total of n= 9 for each combination). Error bars, SD. Groups indicated by the same letter are not statistically different according to ANOVA with Tukey’s post hoc test.

We next asked if MSL10 physically interacts with SYT5 or SYT7. Although SYT5 and SYT7 were not detected in the MSL10 interactome (**Figure 1**), those experiments were performed in seedlings, whereas the suppression of *msl10-3G* phenotypes by *sdm26* and *sdm34* alleles was observed in adult plants. In the mbSUS assay, yeast expressing SYT5 and SYT7 did not grow on minimal media when mated to yeast expressing MSL10 (**Figure 7c**). A FRET-FLIM assay also failed to provide evidence for a direct interaction between MSL10 and SYT proteins, as co-expression of mRFP-labelled SYT5, SYT7, and SYT1 did not shift the fluorescence lifetime of MSL10-GFP (**Figure 7d).** The lack of evidence for physical interactions between MSL10 and SYT1, SYT5, and SYT7 suggests that the observed suppression of the *msl10-3G* phenotype in *sdm26* or *sdm34* mutants is executed indirectly, perhaps through signaling intermediates.

## DISCUSSION

The mechanosensitive ion channel MSL10 has been well studied using electrophysiological approaches (Haswell et al., 2008; Maksaev and Haswell, 2012; Maksaev et al., 2018). Genetic analyses have attributed a variety of roles to MSL10, like the induction of Ca^2+^ transients, reactive oxygen species accumulation, enhanced immune responses, and programmed cell death (Basu and Haswell, 2020; Moe-Lange et al., 2021; Basu et al., 2021), but we lack a clear understanding of how MSL10 activation leads to these downstream signaling outcomes. Studies using multiple gain-of-function *MSL10* alleles found that MSL10 signaling can trigger cell death independently of ion flux (Veley et al., 2014; Zou et al., 2016; Maksaev et al., 2018; Basu et al., 2020), though it remains unknown how this occurs. To advance our understanding of the signaling function of MSL10, we used a combination of genetic, proteomic, and cell biological approaches in an attempt to identify MSL10’s signaling partners. We discovered previously unknown interactions between MSL10, which is localized to the plasma membrane, and proteins in the VAP27 and SYT families, which are integral ER membrane proteins. **Figure 8** outlines these results and provides a framework for the discussion below. We propose a model wherein 1) MSL10’s direct interaction with VAP27s creates EPCSs which 2) has implications for MSL10 function and 3) SYTs and MSL10 interact indirectly to modulate MSL10 signaling and SYT1 localization.

**Figure 8.**
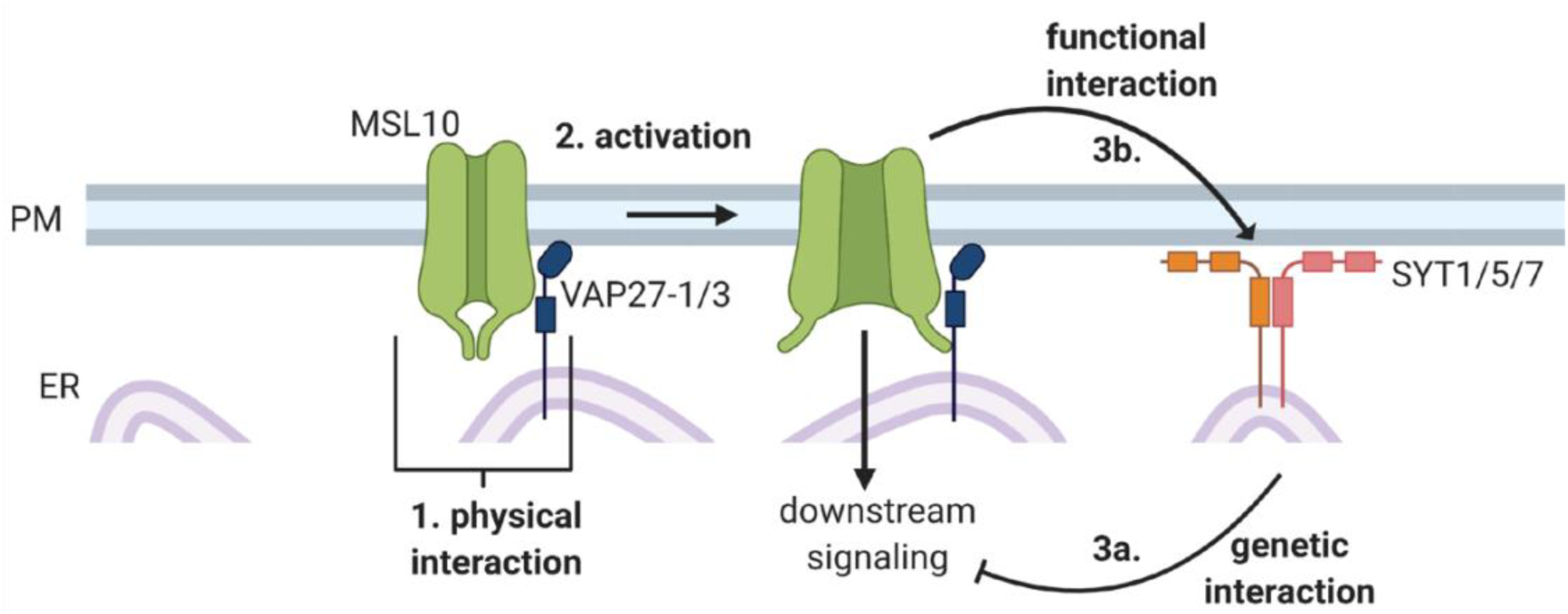
Conceptual model of interactions between MSL10 and EPCS proteins.

### 1. MSL10 physically associates with EPCS proteins

The first indication that MSL10 was part of a protein complex at EPCSs came from our search for proteins that co-immunoprecipitated with MSL10-GFP from seedling microsome extracts. VAP27-1, VAP27-3, and SYT1 were among the most enriched proteins in these pulldowns **(Figure 1)**. Subsequent mbSUS and FRET-FLIM assays support a direct interaction between MSL10 and VAP27-1 and VAP27-3, but not with SYT1 or 11 other proteins tested (**Figure 2**, **Figure 3**). SYT1, ACT8, and AT3G62360 have been detected in other EPCS proteomes (Ishikawa et al., 2020; Kriechbaumer et al., 2015), and were likely found in the MSL10 interactome because of their proximity to VAP27-1 and VAP27-3. Plant EPCSs typically contain either SYT1 or VAP27-1, but SYT1- and VAP27-1-EPCSs are often found adjacent to each other (Siao et al., 2016), suggesting a physical link between two types of EPCSs.

### 2. Implications of the VAP27-1/3 interaction for MSL10 cell death signaling

The only components of our proteome (among 14 tested proteins) that interacted directly with MSL10 were VAP27-1 and VAP27-3 (**Figure 8, point 1**). Broadly speaking, VAPs serve to recruit other proteins or whole protein complexes to the ER membrane. If the client protein is embedded in another organellar membrane, this interaction by definition leads to the formation of a membrane contact site (James and Kehlenbach, 2021). VAP27-1 interacts with SEIPIN2 and SEIPIN3 at ER-lipid droplet contact sites (Greer et al., 2020) and VAP27-3 recruits soluble oxysterol-binding protein-related protein ORP3a to the ER (Saravanan et al., 2009). At EPCSs, Arabidopsis VAP27-1 and VAP27-3 interact with clathrin and are required for normal rates of endocytosis, perhaps by recruiting clathrin to the PM (Stefano et al., 2018). Other VAP27-1 interactors include PM intrinsic protein (PIP)2;5, an aquaporin (Fox et al., 2020), AtEH1/Pan1, a protein that recruits endocytic proteins to autophagosomes that form at VAP27-1-containing EPCSs (Wang et al., 2019), and the actin-binding protein NETWORKED 3C (Wang et al., 2014). The cytosolic domains of VAP27-1 and VAP27-3 can interact with phospholipids (Stefano et al., 2018), which raises the possibility they may not need to interact with a protein in another membrane to create a membrane contact site.

Here we add another VAP27 interactor, one that is associated with mechanical signaling. MSL10 signaling is hypothesized to be activated by membrane tension-induced conformational changes that lead to its dephosphorylation and the activation of its signaling function (Basu et al., 2020). One could imagine that such post-translational modifications disrupt the ability of MSL10 to interact with VAP27-1 and VAP27-3, thereby activating downstream responses.

However, the fact that phosphomimetic (MSL10^7D^), phosphodead (MSL10^7A^), and gain-of-function *msl10-3G* (MSL10 S640L) versions all interacted with VAP27-1 and VAP27-3 (**Figure 2-supplemental figure 1**) implies that MSL10 signaling activation is independent of VAP binding. Rather, MSL10 and VAP27s are likely to interact constitutively, as they interacted both in adult leaves, a tissue type in which MSL10-GFP overexpression promotes cell death signaling (Veley et al., 2014) and in seedlings, a stage where MSL10-GFP overexpression has no effect under normal conditions (Basu and Haswell, 2020).

MSL10 channel and cell death signaling activities are separable (Veley et al., 2014; Maksaev et al., 2018), and VAP27-1 or VAP27-3 could influence either or both of these functions (**Figure 8, point 2**). In *Zea mays*, interaction with VAP27-1 increases the ability of the PM-localized aquaporin *Zm*PIP2;5 to transport water (Fox et al., 2020). Conversely, the mammalian Kv2.1 K^+^ channel forms non-conducting clusters when it interacts with the VAP27-1 homologs VAPA and VAPB (O’Connell et al., 2010; Fox et al., 2013; Johnson et al., 2018). It will be interesting to test if association with VAP27-1 or VAP27-3 alters the channel properties of MSL10, such as its tension sensitivity. Alternatively, interaction with VAP27s could bring ER-localized regulators of MSL10 signaling into proximity, as is the case for an ER-bound phosphatase and its PM receptor substrate (Haj et al., 2012).

### 3A. Point mutations in SYT5 and SYT7 suppress MSL10 signaling

The *msl10-3G* suppressor screen produced two dominant extragenic *sdm* mutants that were successfully mapped to *SYT5* and *SYT7* genes (**Figures 4, 5**). Plant synaptotagmins and homologous proteins in mammals (extended-synaptotagmins, E-SYTs) and yeast (tricalbins) directly bridge the ER and PM via the interaction of their C2 domains with PM phospholipids (Schulz and Creutz, 2004; Min et al., 2007; Giordano et al., 2013; Schapire et al., 2008; Perez-Sancho et al., 2015; Ruiz-Lopez et al., 2021). E-SYTs and tricalbins non-selectively transport glycerolipids between membranes through their synaptotagmin-like mitochondrial lipid-binding (SMP) domains, and Arabidopsis SYT1 and SYT3 are hypothesized to transfer diacylglycerol from the PM to the ER during stress conditions (Ruiz-Lopez et al., 2021). The SYT5 S66F mutation (*sdm26* allele) occurs just outside of the predicted SMP domain of SYT5, and the SYT7 G427R mutation (*sdm34* allele) is found between two predicted C2 domains and near a coiled-coil domain (**Figure 5**). However, both *sdm* alleles were dominant, and both had the same effect of suppressing *msl10-3G* signaling (**Figure 8, point 3a**). Perhaps these lesions, which are in linker regions, influence the large-scale conformational changes that SYTs and E-SYTs are thought to undergo in the presence of Ca^2+^ and certain PM phosphatidylinositol phosphates (Bian et al., 2018; Benavente et al., 2021). This could affect the distance between the ER and PM and the transport of lipids between them, creating a novel lipid environment around MSL10 that might attenuate its ability to activate cell death signaling. Alternatively, the *sdm* mutations in SYT5 and SYT7 might alter the stoichiometry of other proteins at EPCSs, and in turn affect MSL10 function. To test these ideas, lipid transport, phospholipid binding, and interacting proteins should be compared between WT and mutant versions of SYT5 and SYT7.

### 3B. SYT1-EPCSs are expanded in *msl10-3G* plants

EPCSs in plant epidermal cells expand in response to environmental perturbations like cold and ionic stress (Lee et al., 2019, 2020; Ruiz-Lopez et al., 2021). We did not find a role for MSL10 in salinity or mannitol-induced EPCS expansion, nor in the shrinking observed after hypo-osmotic shock (**Figure 3-supplemental figure 1**). However, we did find that SYT1 EPCSs were constitutively expanded in leaf epidermal cells of adult *msl10-3G* plants (**Figure 3**). We did not observe expanded SYT5- or SYT7-EPCSs in *msl10-3G* plants (**Figure 7**). Although SYT1, SYT5, and SYT7 can interact with each other in immunoprecipitations of whole seedling extracts and in bimolecular fluorescence complementation assays (Ishikawa et al., 2020; Lee et al., 2020), perhaps they are not in a complex together in all cell types or developmental stages.

Why are SYT1-EPCSs expanded in *msl10-3G* leaves? We previously reported that the *msl10-3G* allele promotes a stronger cytosolic Ca^2+^ transient in response to hypo-osmotic cell swelling than is seen in WT seedlings (Basu and Haswell, 2020). The affinity of SYT1 for PM phospholipids is partially dependent on Ca^2+^ (Schapire et al., 2008; Perez-Sancho et al., 2015), suggesting that MSL10 could affect SYT1 function. Alternatively, perhaps EPCSs are expanded in *msl10-3G* cells because these cells are already ‘stressed’; *msl10-3G* plants constitutively express markers of wounding and abiotic stress (Zou et al., 2016; Basu et al., 2020). If overactive stress responses in *msl10-3G* plants increase PM phosphatidylinositol 4,5-bisphosphate (PI(4,5)P2) levels, as wounding (Mosblech et al., 2008) or saline conditions (Lee et al., 2019) do, SYT1-EPCS expansion could be promoted. Both of these scenarios are consistent with the fact that we do not observe altered EPCSs in null *msl10-1* leaves. At the moment, the effects we observe on SYT1 area are limited to the gain-of-function *msl10-3G* allele.

However, we did find genetic interactions between the null *msl10-1* allele and a SYT1-GFP overexpression transgene (**Figure 3-Supplementary Table 1**). In addition, we were unable to isolate any adult plants overexpressing VAP27-3-GFP in either the null *msl10-1* or gain-of-function *msl10-3G* lines. Taken together, these unexpected genetic results may indicate that the stoichiometry of proteins at plant EPCSs is tightly balanced, and that when disturbed, perturbations of components even in opposing directions can be detrimental. In support of this idea, VAP27-1 gain-of-function and loss-of-function lines both have abnormal root hairs (Wang et al., 2016). Transient overexpression of two EPCS proteins at the same time can drastically alter plant ER and EPCS morphology or even cause necrosis (Wang et al., 2016; Ruiz-Lopez et al., 2021). Additionally, a yeast strain missing all EPCS tethering proteins is viable but cannot tolerate the loss of *OSH4,* a redundant lipid-transport protein (Quon et al., 2018, 2022). Thus, we interpret the synthetic lethality of *MSL10* alleles and VAP27-3 or SYT1 overexpression transgenes as additional evidence that MSL10 functions at plant EPCSs, and we speculate that the ectopic cell death observed in plants overexpressing MSL10-GFP (Veley et al., 2014; Basu et al., 2020) may be a consequence of altered stoichiometry of EPCS proteins and/or dysfunction of EPCSs. Future studies should examine the dynamics of MSL10, SYTs, and VAP27s in the presence, absence, and overexpression of each other—similar to the study of (Siao et al., 2016)—to begin to understand the influence they have on each other.

### Implications of having a mechanosensitive ion channel at EPCSs

To our knowledge, MSL10 is the first mechanosensitive ion channel to be found at plant or animal EPCSs, but this may be an unsurprising location to find a mechanosensory protein in any system. It is hypothesized that plant EPCSs interact indirectly with the cell wall (Wang et al., 2017). VAP27-1 and SYT1 are found at Hechtian strands (Wang et al., 2016; Lee et al., 2020), sites of connection between the PM and the cell wall, and the mobility of VAP27-1 is constrained by the presence of a cell wall (Wang et al., 2016). Additionally, plant EPCSs link to the actin and microtubule cytoskeletons (Wang et al., 2014; Zang et al., 2021), which might convey or transduce mechanical information to or from the ER-PM-cell wall interface. By placing the mechanosensitive ion channel MSL10 at EPCSs, our results indicate that EPCSs will be an important nexus for understanding plant mechanotransduction cascades in a cellular context.

## MATERIALS AND METHODS

### Plant lines and growth conditions

All *Arabidopsis thaliana* lines used in this study are in the Col-0 ecotype. *msl10-3G (rea1)* seeds were derived from an ethyl methanesulfonate (EMS) mutant screen (Zou et al., 2016) and subsequently backcrossed twice (once to parental *RAP2.6::Luc* background and once to Col-0) to remove additional EMS-induced mutations. T-DNA insertion mutants *syt1* (SAIL_775_A08), *syt5* (SALK_03961), and *syt7* (SALK_006298) (Ishikawa et al., 2020) and *msl10-1* (Haswell et al., 2008) were obtained from the Arabidopsis Biological Resource Center. *UBQ:MAPPER-GFP* seeds were a gift from Abel Rosado (Lee et al., 2019). Unless otherwise specified, plants were grown on soil at 22°C under a constant light regime (120 µmol m^-2^ s^-1^).

### Genotyping

DNA was isolated by homogenizing tissue in 300 µL crude extraction buffer (200 mM Tris-HCl pH 7.5, 250 mM NaCl, 250 mM EDTA, and 0.5% sodium dodecyl sulfate) followed by precipitation with an equal volume of isopropanol. Mutant lines were genotyped using the primers indicated in Table S1. The *msl10-3G* point mutation was genotyped using primers 663 and 702 followed by digestion with the *Taq1* restriction enzyme, which cuts only the WT *MSL10* allele. The *sdm26 (SYT5 S66F)* point mutation was genotyped using primers 4155 and 4156 followed by digestion with the *Taq1* restriction enzyme, which cuts the mutant, but not WT *SYT5* sequence. The *sdm34 (SYT7 G427R)* point mutation was genotyped using dCAPs primers 4231 and 4232 and digestion with the *DdeI* enzyme, which cuts the mutant but not the WT *SYT7* allele

### Cloning and generation of transgenic plants

To make *SYT5g S66F* and *SYT7g G427R* constructs, the *SYT5* and *SYT7* genomic sequences were amplified from pGWB553 SYT5g-mRFP and pGWB553 SYT7g-mRFP vectors (Ishikawa et al., 2020) which were a gift from Kazuya Ishikawa and cloned into the pENTR vector using the pENTR/D-TOPO Cloning Kit (Thermo Fisher). These pENTR constructs were used as templates for site-directed mutagenesis to introduce *SYT5 S66F* or *SYT7 G427R* mutations (primers in Table S1) The mutated genomic sequences were subcloned back into pGWB553 vectors using Gibson assembly with NEBuilder Hifi DNA Assembly Master Mix (NEB). The WT constructs included a C-terminal mRFP tag (Ishikawa et al., 2020), and the *sdm* constructs had a short, 31aa tag before an early stop codon was reached. The resulting constructs were transformed into *msl10-3G* plants using *Agrobacterium tumefaciens* GV3101 and the floral dip method (Clough and Bent, 1998). T1 individuals were identified based on hygromycin resistance.

To make *UBQ:SYT1-GFP, UBQ:SYT5-GFP, UBQ:SYT7-GFP, UBQ:VAP27-1-GFP,* and *UBQ:VAP27-3-GFP* constructs, the *SYT1, SYT5, SYT7, VAP27-1,* and *VAP27-3* coding sequences were amplified from Col-0 cDNA using primers in Table S1 and cloned into pENTR using pENTR/D-TOPO, then subcloned into the pUBC-GFP-DEST vector (Grefen et al., 2010) using LR Clonase II (Thermo Fisher recombination. The resulting constructs were transformed introduced into Col-0 plants and transformed individuals were identified based on Basta resistance. T2 plants with moderate GFP fluorescence were crossed to *msl10-1* and *msl10-3G* plants, and homozygous F2 siblings were identified by genotyping and by screening for Basta resistance. To make *UBQ:mRFP-VAP27-1, UBQ:mRFP-VAP27-3, UBQ:SYT1-mRFP, UBQ:SYT5-mRFP, UBQ:SYT7-mRFP,* and *UBQ:MSL10-GFP,* Clonase II recombination was used to subclone the coding sequences of *VAP27-1* and *VAP27-3* from pENTR into the pUBN-RFP-DEST vector, *SYT1*, *SYT5*, and *SYT7* into pUBC-RFP-DEST, and *MSL10* into pUBC-GFP-DEST (Grefen et al., 2010).

To make *pK7-mRFP-VAP27-3g*, the *VAP27-3* genomic sequences were amplified from Col-0 genomic DNA. Using Gibson assembly, this was cloned into the *pK7FWG2* vector backbone, deleting the GFP tag and adding an N-terminal mRFP tag. For co-localization studies, this construct was transformed into Col-0 plants expressing a *MSL10p:MSL10-GFP* transgene (Haswell et al., 2008). T1 plants were identified by kanamycin resistance.

### Microsome isolation and immunoprecipitation

Seeds of Col-0 and *35S:MSL10-GFP* (line 12-3, (Veley et al., 2014; Basu et al., 2020b)) were densely sown on 1X Murashige and Skoog (MS) plates supplemented with 3% sucrose and grown vertically for 7 days in a 16 hr light/8 hr dark regime. Seedlings (1 g per replicate) were flash frozen in liquid nitrogen and homogenized to a fine powder using a mortar and pestle. Protein extraction and microsome isolation protocols were modified from (Abas and Luschnig, 2010). 1.5 mL of extraction buffer (100 mM Tris-HCl pH 7.5, 25% sucrose, 5% glycerol, 3.3% polyvinylpyrolidone, 10 mM EDTA, 10 mM EGTA, 5 mM KCl, 1 mM DTT, 0.1 mM PMSF, 2 µM leupeptin, 1 µM pepstatin, 1X plant protease inhibitor cocktail (Sigma P9599), and 1X phosphatase inhibitor cocktails 2 (Sigma P5726) and 3 (Sigma P0044)) was added directly to the mortar and samples were homogenized in buffer for 2 min, then transferred to 1.5 mL tubes and incubated on ice for 10 min. Homogenates were centrifuged at 600g for 3 min (1 replicate) or 10,000g for 10 min (3 replicates) at 4°C to pellet cell debris and organelles. The supernatant was transferred to fresh tubes on ice, and the pellets were resuspended in half of the initial volume of extraction buffer, using small plastic pestles. Resuspensions were centrifuged as above. Pooled supernatants were diluted 1:1 with ddH_2_0, then divided among 1.5 mL tubes, each with a maximum volume of 200 mL. Microsomes were pelleted by centrifugation at 21,000g for 2 hr at 4°C, and the supernatant was discarded.

Microsomal pellets were then resuspended in a total volume of 0.5 mL solubilization buffer (20 mM Tris-HCl pH 7.5, 150 mM NaCl, 2 mM EDTA, 10% glycerol, 0.5% Triton X-100, 0.25% NP-40, 0.1 mM PMSF, 2 µM leupeptin, 1 µM pepstatin, 1X plant protease inhibitor cocktail, and 1X phosphatase inhibitor cocktails 2 and 3) using small plastic pestles. Resuspended microsomes were incubated with end-over-end rotation at 4°C for 1 hr. Meanwhile, 65 µL of GFP-Trap Magnetic Agarose beads (Chromotek) per sample was prepared by washing twice with 1 mL 10 mM Tris-HCl, 150 mM NaCl, 0.5 mM EDTA. To this was added 400 µL of solubilized microsomes and 100 µL of solubilization buffer. Proteins were immunoprecipitated overnight with end-over-end rotation at 4°C. Beads were collected with a magnetic rack, and the flow-through was discarded. Beads were washed 3 times with 1 mL IP Wash Buffer 1 (20 mM Tris-HCl pH 7.5, 150 mM NaCl, 10% glycerol, 2 mM EDTA, 1% Triton X-100, and 0.5% NP-40), then 6 times with IP Wash Buffer 2 (20 mM Tris-HCl pH 7.5, 150 mM NaCl, 10% glycerol, 2 mM EDTA), switching to fresh tubes every other wash.

### Tandem mass spectrometry

Proteins were eluted from the GFP-Trap beads by adding 100 µL of 8 M urea, then reduced in 10 mM dithiothreitol for 1 hr at RT, and alkylated in the dark (50 mM 2-iodoacetamide) for 1 hr at RT. Excess alkylating agent was quenched with 50 mM dithiothreitol for 5 min at RT. Samples were diluted with 900 µl of 25 mM ammonium bicarbonate and digested overnight at 37°C in the presence of 0.35 µg of sequencing grade modified porcine trypsin (Promega). Peptides were vacuum-dried in a centrifugal evaporator to approximately 250 µl, acidified with 10% trifluoroacetic acid (TFA) (pH<3), desalted and concentrated on a 100 µl Bond Elut™ OMIX C18 pipette tip (Agilent Technologies A57003100) according to the manufacturer’s instructions.

Peptides were eluted in 50 µl of 75% aceonitrile, 0.1% acetic acid, vacuum dried in a centrifugal evaporator (Savant Instruments, model number SUC100H), and resuspended in 17 µl 5% acetonitrile, 0.1% formic acid.

Nano-scale liquid chromatography (LC) separation of tryptic peptides was performed on a Dionex Ultimate™ 3000 Rapid Separation LC system (Thermo Fisher). The protein digests were loaded onto a 20 μl nanoViper sample loop (Thermo Fisher), and separated on a C18 analytical column (Acclaim PepMap RSLC C18 column, 2 μm particle size, 100 Å pore size, 75 µm x 25 cm (Thermo Fisher)) by the application of a linear 2 hr gradient from 4% to 36% acetonitrile in 0.1% formic acid, with a column flow rate set to 250 nL/min. Analysis of the eluted tryptic peptides was performed online using a Q Exactive™ Plus mass spectrometer (Thermo Scientific) possessing a Nanospray Flex™ Ion source (Thermo Fisher) fitted with a stainless steel nano-bore emitter operated in positive electro-spray ionisation (ESI) mode at a capillary voltage of 1.9 kV. Data-dependent acquisition of full MS scans within a mass range of 380-1500 m/z at a resolution of 70,000 was performed, with the automatic gain control (AGC) target set to 3.0 x 10^6^, and the maximum fill time set to 200 ms. High energy collision-induced dissociation (HCD) fragmentation of the top 8 most intense peaks was performed with a normalized collision energy of 28, with an intensity threshold of 4.0 x 10^4^ counts and an isolation window of 3.0 m/z, excluding precursors that had an unassigned, +1 or >+7, charge state. MS/MS scans were conducted at a resolution of 17,500, with an AGC target of 2 x 10^5^ and a maximum fill time of 300 ms. Dynamic exclusion was performed with a repeat count of 2 and an exclusion duration of 30sec, while the minimum MS ion count for triggering MS/MS was set to 4 x 10^4^ counts. The resulting MS/MS spectra were analyzed using Proteome Discoverer™ software (version 2.0.0.802, Thermo Fisher), which was set up to search the *Arabidopsis thaliana* proteome database, as downloaded from www.tair.com (TAIR10_pep_20101214). Peptides were assigned using SEQUEST HT (Eng et al., 1994), with search parameters set to assume the digestion enzyme trypsin with a maximum of 1 missed cleavage, a minimum peptide length of 6, precursor mass tolerances of 10 ppm, and fragment mass tolerances of 0.02 Da. Carbamidomethylation of cysteine was specified as a static modification, while oxidation of methionine and N-terminal acetylation were specified as dynamic modifications. The target false discovery rate (FDR) of 0.01 (strict) was used as validation for peptide-spectral matches (PSMs) and peptides. Proteins that contained similar peptides and which could not be differentiated based on the MS/MS analysis alone were grouped, to satisfy the principles of parsimony. Label-free quantification as previously described (Silva et al., 2006) was performed in Proteome Discoverer™ with a minimum Quan value threshold of 0.0001 using unique peptides, and “3 Top N” peptides used for area calculation. All samples were injected in duplicate, and the resulting values were averaged. The mass spectrometry proteomics data have been deposited to the ProteomeXchange Consortium via the PRIDE partner repository (Perez-Riverol et al., 2019) with the dataset identifier PXD018747.

Using the Perseus platform (Tyanova et al., 2016), intensity values from mass spectrometry were log_2_ imputed and missing values were replaced with random numbers from a Gaussian distribution with a width of 0.3 and a downshift of 1.8. Statistical significance was determined using t-tests. Only proteins with > 8 peptide spectrum matches were included in volcano plots.

### Mating-based split ubiquitin (mbSUS) assay

The coding sequence for the 14 proteins selected from the MSL10 interactome were amplified from Col-0 cDNA using primers in Table S1 and cloned into *pENTR* using pENTR/D-TOPO, then subcloned into the *pK7FWG2* destination vector (Karimi et al., 2002) or BiFC destination vectors (Gehl et al., 2009) using LR Clonase II recombination. These constructs were used as templates for PCR amplification with attB1 For and attB2 Rev primers (Table S1). Following the protocol of (Obrdlik et al., 2004; Basu et al., 2020b), attB-flanked inserts were combined with linearized vectors and transformed into yeast for recombinational *in vivo* cloning. Inserts were cloned into *pMetYCgate* for a C-terminal fusion with Cub, *pXNgate21-3HA* for a C-terminal fusion with NubG, or *pNXgate33-3HA* for an N-terminal NubG fusion. For integral membrane proteins split-ubiquitin tags were predicted to lie in the cytosol. For soluble proteins, the NubG tag was placed on the terminus where fusions had previously reported to be tolerated (or, for unstudied proteins, where homologous proteins had been tagged). NubG vectors and inserts were transformed into THY.AP5 cells and selected on Synthetic Complete (SC) plates lacking tryptophan and uracil. Cub vectors and inserts were transformed into THY.AP4 cells and selected on SC plates lacking leucine. Transformed cells were mated and diploids selected on SC media lacking tryptophan, uracil, and leucine. Overnight cultures of diploid cells were pelleted, resuspended in dH_2_0 to an OD_600_ of 1.0, and 4 µL of a 10X dilution series were spotted onto Synthetic Minimal (SD) or SC+Ade+His media. Growth was assessed 3 days after plating; growth on SC+Ade+His media tested the presence of both constructs. To quantify the strength of interactions, β-galactosidase activity in liquid cultures was assayed using CPRG as substrate as described in the Yeast Protocols Handbook (Takara).

## FRET-FLIM

*UBQ:mRFP-VAP27-1, UBQ:mRFP-VAP27-3, UBQ:SYT1-mRFP, UBQ:SYT5-mRFP, UBQ:SYT7-mRFP,* and *UBQ:MSL10-GFP* plasmids were transformed into *A. tumefaciens* GV3101. Following the protocol of (Waadt and Kudla, 2008), construct pairs were co-infiltrated into *Nicotiana benthamiana* leaves along with *A. tumefaciens* strain AGL-1, which harbors p19 to suppress gene silencing. 5 days post-infiltration, leaves were imaged using a Leica TCS SP8 Multiphoton microscope fitted with an HC PL IRAPO 40x/1.10 WATER objective. The tunable multiphoton laser was adjusted to its optimum excitation for EGFP (920 nm), and fluorescence lifetimes were recorded in an emission range of 595-570 nm. Using the Leica LASX software’s FLIM tool, an n-Exponential Reconvolution model with one component was used to calculate the average fluorescence lifetime of GFP per image.

### Co-localization analysis

Leaves of plants co-expressing *MSL10p:MSL10-GFP* and *mRFP-VAP27-3g* were imaged using an Olympus FV3000 confocal microscope with a UPLSAPO 100XS oil-immersion objective. 8 to 12 Z-slices were captured at the equator of abaxial leaf epidermal cells, and these Z-stacks were deconvolved. For each image, ROIs were defined at the periphery of 4 different cells. Co-localization was quantified using the ‘Co-localization’ tool of the Olympus cellSens software, using the ‘Rectangle’ mode to automatically estimate thresholds, and the mean of the Mander’s coefficients was calculated from the 4 ROIs in 4 Z-slices.

### Confocal microscopy and quantification of ER-plasma membrane contact sites

Lines expressing MAPPER-GFP, SYT1-GFP, SYT5-GFP, SYT7-GFP, VAP27-1-GFP, and VAP27-3-GFP under the control of the *UBQ10* promoter were visualized using an Olympus FV3000 confocal microscope with a UPLSAPO 100XS oil-immersion objective. GFP was excited using a 488nm laser and detected in the 500-540nm range. Chlorophyll autofluorescence was excited by the same laser and detected in the 650-750nm range. Z-stacks were taken of abaxial leaf epidermal cells beginning at the top of the cell and ending with an equatorial slice. Z-stacks were deconvolved with the Olympus CellSens software using the Advanced Maximum Likelihood Algorithm with 5 iterations. The area of MAPPER-GFP or SYT1-GFP puncta were quantified using Fiji (Schindelin et al., 2012). Deconvolved Z-stacks were converted to a Z-projection (sum slices for MAPPER-GFP and maximum intensity for SYT1-GFP) and the area of each cell was traced and set as an ROI, excluding the periphery of cells where puncta were typically overlapping. After thresholding (between 25-255 for MAPPER-GFP and 100-255 for SYT1-GFP, the ‘Analyze Particles’ function was used to quantify the % of cell area that the puncta represented for each ROI.

### Identification of suppressed death from msl10-3G (sdm) mutants

250 mg of backcrossed *msl10-3G* seeds (approximately 12,500 seeds) were treated with 0.4% ethyl methanesulfonate (EMS) as described in (Kim et al., 2006). Mutagenized seeds were sown directly on soil in 40 pools, stratified for 2 days at 4°C, then transferred to a 22°C growth chamber. *sdm* mutants were identified based on increased height compared to parental *msl10-3G* plants 4-5 weeks after sowing, each from individual pools. When multiple plants with *sdm* phenotypes were seen in the same M2 pool they were assumed to be from the same parent. *sdm* mutants were genotyped to ensure they had the *msl10-3G* point mutation. To see if *sdm* mutants harbored second-site mutations in the *MSL10* gene, the locus was PCR amplified using primers 3781 and 3782 and Sanger-sequenced using primers 663, 699, 701, 1611, 2227, and 3789 (Table S1).

*sdm26 and sdm34* were backcrossed to *msl10-3G* plants, and rosette leaves from 30-50 F2 progeny were separated into two pools based on phenotype: *msl10-3G* (dwarfed) or *sdm* (suppressed). Genomic DNA was extracted from pooled tissue following the protocol described in (Thole et al., 2014) and submitted to the Genome Technology Access Center at the McDonnell Genome Institute (GTAC@MGI) at the WUSTL Medical Center. Libraries were prepared using the Kapa HyperPrep Kit PCR-free (Roche) and sequenced on an Illumina NovaSeq 6000 S4 Flowcell using 150 nt paired-end reads and 80X coverage. GTAC@MGI aligned reads to the *Arabidopsis thaliana* Col-0 reference genome (TAIR10.1 assembly), called variants using SAMtools (Li et al., 2009), and annotated them using snpEff (Cingolani et al., 2014). Variants were filtered to include those with a quality score of >20 and a total depth of >5. SNPs that were present in multiple *sdm* mutants were removed, as they were likely present in the parental *msl10-3G* line. For each of the retained SNPs, the allele frequency (mutant/reference) was plotted against chromosomal position.

### Alignment of SYT5 and SYT7 protein sequences

SYT5 and SYT7 homologs in other plant species were identified using the BLAST tools in Phytozome 13 or NCBI using the Arabidopsis SYT5 and SYT7 amino acid sequences as queries. To remove sequences that were orthologous to other Arabidopsis synaptotagmins, we aligned the obtained sequences to the protein sequences of the 7 known synaptotagmins in Arabidopsis and constructed a Neighbor-Joining phylogenic tree in Mega 11. We then considered only those sequences that were in the same clade as *At*SYT5 or *At*SYT7 to be SYT5 or SYT7 homologs. SYT5 homologs identified with this method and shown in Figure 5c have the following accession numbers from Phytozome: *B. rapa* B.rapaFPsc v1.3|Brara.J00373.1.p, *V. Vinifera* v2.1|VIT_211s0118g00230.2, *P. trichocarpa* v4.1|Potri.018G025000.3.p, *O. sativa* v7.0|LOC_Os04g55220.1, *B. distachyon* v3.2|Bradi5g23880.2.p. From NCBI: *N. tabacum* XP_016446163.1. SYT7 homologs identified in Phytozome include *B. rapa* B.rapaFPsc v1.3|Brara.D00127.1.p, *V. vinifera* v2.1|VIT_215s0048g01410.1, *P. trichocarpa* v4.1|Potri.014G072800.2.p, *O. sativa* v7.0|LOC_Os07g22640.1, *B. distachyon* v3.2|Bradi1g52680.1.p. From NCBI: *N. tabacum* XP_016486625.1.

### Immunoblotting

Rosette leaves were flash frozen and homogenized in a microcentrifuge tube using a small plastic pestle. 4 µL of 2X sample buffer was added for every 1 mg of tissue, then this mixture was denatured for 10 min at 70°C and cell debris pelleted by centrifugation at 5000g for 1 min. Supernatants were resolved on 10% SDS-PAGE gels and transferred overnight to PVDF membranes (BioRad) at 100 mA. Blocking and antibody incubations were performed in 5% non-fat dry milk in 1X TBS-T buffer. MSL10 tagged with GFP was detected using an anti-GFP antibody (Takara #632380) for 16 hr at a dilution of 1:5000, followed by a 1 hr incubation in HRP-conjugated goat-anti-mouse secondary antibody at a 1:10,000 dilution (Millipore-Sigma #12-349). Blots were stripped and re-probed with anti-α-tubulin (Millipore-Sigma T5168, 1:30,000 dilution) for 1 hr. Proteins were detected using the SuperSignal West Dura Extended Duration Substrate (Thermo Fisher).

### Gene expression analysis

Rosette leaves were flash frozen in liquid nitrogen and homogenized into a powder. RNA was extracted using RNeasy Kit (Qiagen) following the manufacturer’s instructions for plant RNA isolation and on-column DNase digestion. cDNA was synthesized using M-MLV reverse transcriptase (Promega) and oligo(dT) priming. qRT-PCR was performed in technical triplicate using the SYBR Green PCR Master Mix (Thermo Fisher) kit, with primers specific to *SYT5, SYT7,* or *ELONGATION FACTOR 1α (EF1 α)* transcripts (Table S1) on a StepOne Plus Real-time PCR System (Applied Biosystems).

### Accession numbers

The genes utilized in this study have the following Arabidopsis Genome Initiative locus codes: *MSL10 (At5G12080), VAP27-1 (At3G60600), VAP27-3 (At2G45140), SYT1 (At2G20990), SYT5 (At1G05500), SYT7 (At3G61050), ACTIN 8 (ACT8, At1G49240), DYNAMIN-LIKE 1 (DL1, At5G42080), RAB GTPase homolog 1C (RAB1c, At4G17530), METHIONINE OVERACCUMULATOR 3 (MTO3, At3G17390), COATOMER ALPHA-1 SUBUNIT (αCOP1, At1G62020),* unnamed protein with a carbohydrate-binding like fold *(At3G62360),* unnamed protein-M28 Zn-peptidase nicastrin (*At3G44330), RAS-RELATED NUCLEAR PROTEIN 1 (RAN1, At5G20010), CATALASE 2 (CAT2, At4G35090), LOW EXPRESSION OF OSMOTICALLY RESPONSIVE GENES (LOS1, At1G56070), REGULATORY PARTICLE TRIPLE-A 1A (RPT1a, At1g53750), POTASSIUM CHANNEL IN ARABIDOPSIS THALIANA 1 (KAT1, At5G46240)*.

### Statistical analyses

Statistical analyses were performed in R Studio (v4.1.2), except for Chi-squared tests which were performed in Microsoft Excel. Shapiro-Wilk tests were used to test for normality. The *car* and *agricolae* packages were used to perform ANOVAs and indicated post-hoc tests, and *FSA* and *rcompanion* packages for Kruskal-Wallis and Dunn’s post-hoc tests. Data was visualized using R Studio *ggplot2*, GraphPad Prism 7, and Excel. The Venn diagram shown in Figure 1b was created using http://bioinformatics.psb.ugent.be/webtools/Venn/.

## Supporting information

Supplemental Move 3

## ACKNOWLEDGEMENTS

We thank Fionn McLoughlin for performing mass spectrometry experiments and for helpful guidance about immunoprecipitations and data analysis. These experiments were performed at the proteomic facility in the Biology Department at Washington University in St. Louis. Heather Grossman generated the amino acid alignments in Figure 5c. Kazuya Ishikawa (Utsonomiya University) provided *SYT5g-mRFP* and *SYT7g-mRFP* plasmids and shared their full SYT1 immunoprecipitation-mass spectrometry dataset. Abel Rosado (University of British Columbia) provided the *UBQ:MAPPER-GFP* line. We thank the staff of the Jeanette Goldfarb Plant Growth Facility for plant growth assistance. Whole genome sequencing in this publication was made possible in part by Grant Number UL1 RR024992 from the NIH-National Center for Research Resources (NCRR). This work was supported by HHMI-Simons Faculty Scholar Grant 55108530 to E. S. H., National Science Foundation grant MCB 1253103 to E. S. H., and the NSF Center for Engineering Mechanobiology grant CMMI-1548571. J.M.C. was supported by NSF Graduate Research Fellowship DGE-1745038 and a William H. Danforth Plant Sciences Fellowship. Figure 8 was created using BioRender.com.

