## Supplemental Figures for "Unbiased proteomic and forward genetic screens reveal that mechanosensitive ion channel MSL10 functions at ER-plasma membrane contact sites in *Arabidopsis thaliana*"

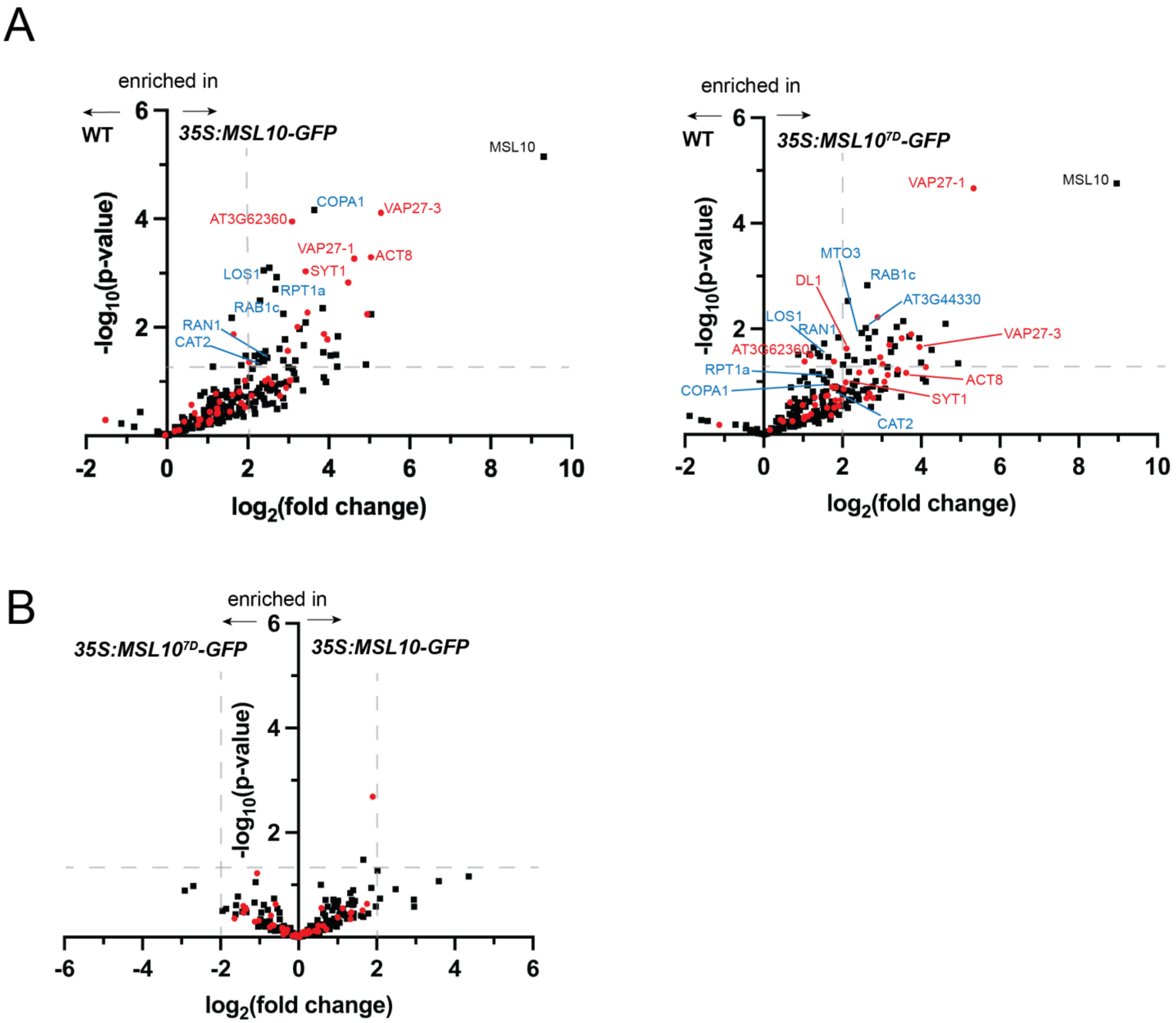


**Figure 1-supplemental figure 1. Similar proteins were identified in MSL10-GFP and MSL107D-GFP immunoprecipitations.** (A) Volcano plots showing the preferential abundance of proteins detected in immunoprecipitations of *35S:MSL10-GFP* (left, reproduced from Figure 1a) or *35S:MSL10$^{7D}$-GFP* (right) compared to those of mock precipitations of WT Col-0 seedlings. (B) displays the relative abundance of proteins in *35S:MSL10-GFP* (right) vs *35S:MSL10$^{7D}$-GFP* (left) immunoprecipitations. Proteins were identified by LC-MS/MS and the average abundance of each was quantified from the MS1 precursor ion intensities, and only those proteins with at least 8 peptide spectral matches are shown. Each protein is plotted based on its -$\log_{10}$(p-value) of significance based on 4 biological replicates relative to its $\log_2$(fold change) of abundance. Data points indicated as red circles have been previously detected in interactomes of SYT1 (Ishikawa et al., 2020), VAP27-1/3 (Stefano et al., 2018), RTNLB3/6 (Kriechbaumer et al., 2015), and VST1 (Ho et al., 2016). Labelled are proteins that were selected for further testing in Figure 2a, selected because in either the MSL10-GFP and/or MSL107D-GFP co-immunoprecipitations they were above the cutoffs indicated as dashed gray lines: a $\log_2$(fold change) > 4 and p-values <0.05. Those with red labels have been found previously in EPCS interactomes, and those in blue have not.

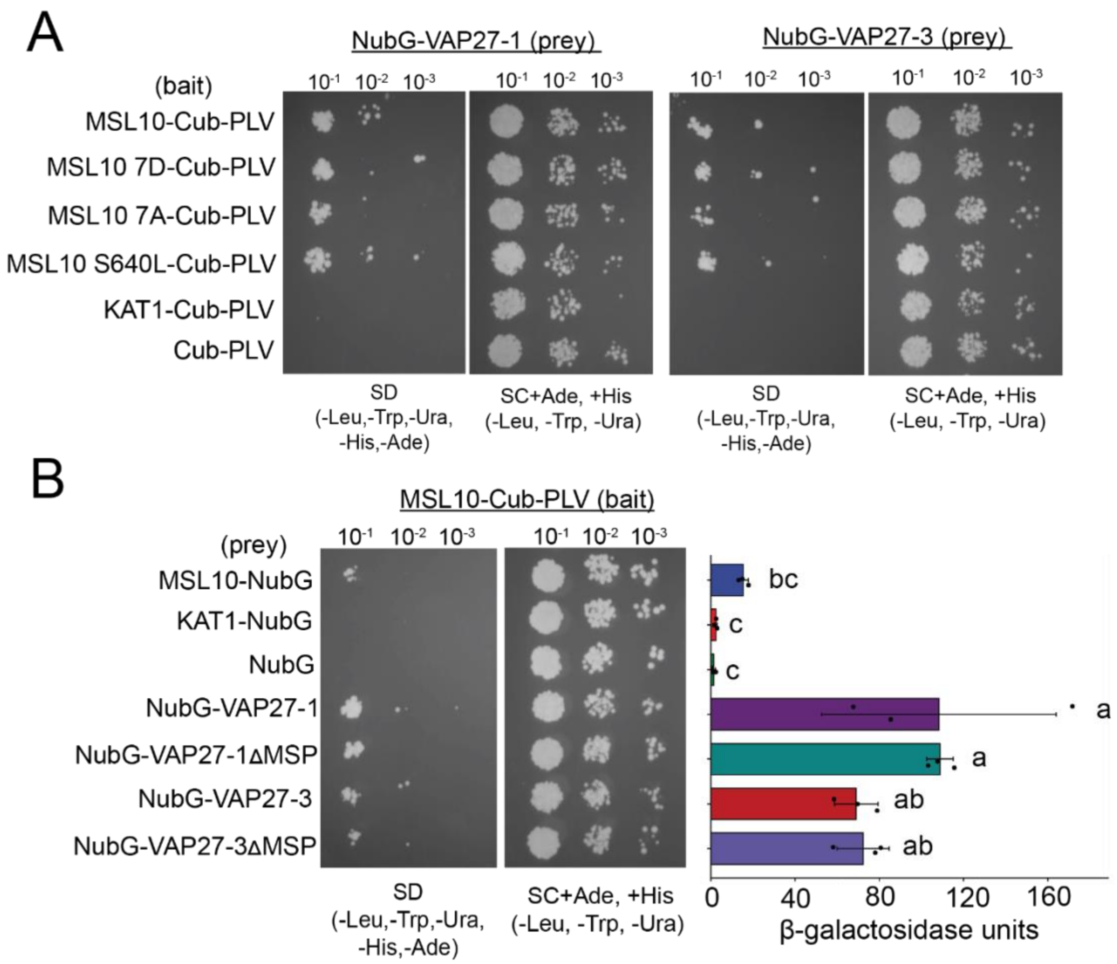


**Figure 2-supplemental figure 1. MSL10 signaling mutants interact with VAP27-1 and VAP27-3, and the VAP27 MSP domain is dispensable for interaction.** Mating-based split-ubiquitin assays testing **(A)** the interaction of full-length VAP27-1 and VAP27-3 mutant versions of full-length MSL10 and **(B)** the interaction of full-length MSL10 with variants of VAP27-1 and VAP27-3 that lacked their major sperm protein (MSP) domain (VAP27-1Δ6-125 and VAP27-3Δ23-142). At right in **(B)**: Measurements of β-galactosidase activity in a liquid-based assay testing the same combinations at left, using CPRG as substrate.

**Figure 3-supplemental table 1. Segregation of *MSL10* alleles in crosses to lines overexpressing GFP-labelled EPCS proteins.** *msl10-1* and *msl10-3G* plants were crossed to lines expressing GFP-labelled VAP27-1, VAP27-3, SYT1, SYT5, or SYT7 under the control of the *UBQ10* promoter. F2 plants (or F3 offspring of heterozygous F2 plants) were selected based on Basta resistance driven by the *UBQ:GFP* transgenes, and resistant plants were genotyped for the indicated *MSL10* alleles. Chi-squared tests were calculated based on a predicted 1:2:1 segregation ratio. Crosses that had significant deviations (p <0.05) from expected ratios are in bold.

| | **# Basta resistant offspring with indicated genotypes** | | | | |
|---|---|---|---|---|---|
| **Parental genotype** | *MSL10* / *MSL10* | *MSL10* / *msl10-3G* | *msl10-3G* / *msl10-3G* | $\boldsymbol{X^2}$ | **p** |
| *UBQ:VAP27-1-GFP* / - ; *MSL10* / *msl10-3G* | 6/25 (24%) | 16/25 (64%) | 3/25 (12%) | 2.68 | 0.26 |
| ***UBQ:VAP27-3-GFP* / - ; *MSL10* / *msl10-3G*** | **12/33 (36%)** | **21/33 (64%)** | **0/33 (0%)** | **11.18** | **0.004** |
| *UBQ:SYT1-GFP* / - ; *MSL10* / *msl10-3G* | 6/21 (29%) | 12/21 (57%) | 3/21 (14%) | 1.29 | 0.53 |
| *UBQ:SYT5-GFP* / - ; *MSL10* / *msl10-3G* | 5/21 (24%) | 7/21 (33%) | 9/21 (43%) | 3.86 | 0.15 |
| *UBQ:SYT7-GFP* / - ; *MSL10* / *msl10-3G* | 9/40 (23%) | 23/40 (57%) | 8/40 (20%) | 0.95 | 0.62 |
| | *MSL10* / *MSL10* | *MSL10* / *msl10-1* | *msl10-1* / *msl10-1* | | |
| *UBQ:VAP27-1-GFP* / - ; *MSL10* / *msl10-1* | 6/28 (21%) | 17/28 (61%) | 5/28 (18%) | 1.36 | 0.51 |
| ***UBQ:VAP27-3-GFP* / - ; *MSL10* / *msl10-1*** | **7/36 (19%)** | **29/36 (81%)** | **0/36 (0%)** | **16.17** | **0.0003** |
| ***UBQ:SYT1-GFP* / - ; *MSL10* / *msl10-1*** | **24/74 (33%)** | **46/74 (62%)** | **4/74 (5%)** | **15.19** | **0.0005** |
| *UBQ:SYT5-GFP* / - ; *MSL10* / *msl10-1* | 7/23 (30%) | 8/23 (35%) | 8/23 (35%) | 2.22 | 0.33 |
| *UBQ:SYT7-GFP* / - ; *MSL10* / *msl10-1* | 16/42 (38%) | 17/42 (41%) | 9/42 (21%) | 3.86 | 0.15 |
| **Expected ratios** | **25%** | **50%** | **25%** | | |

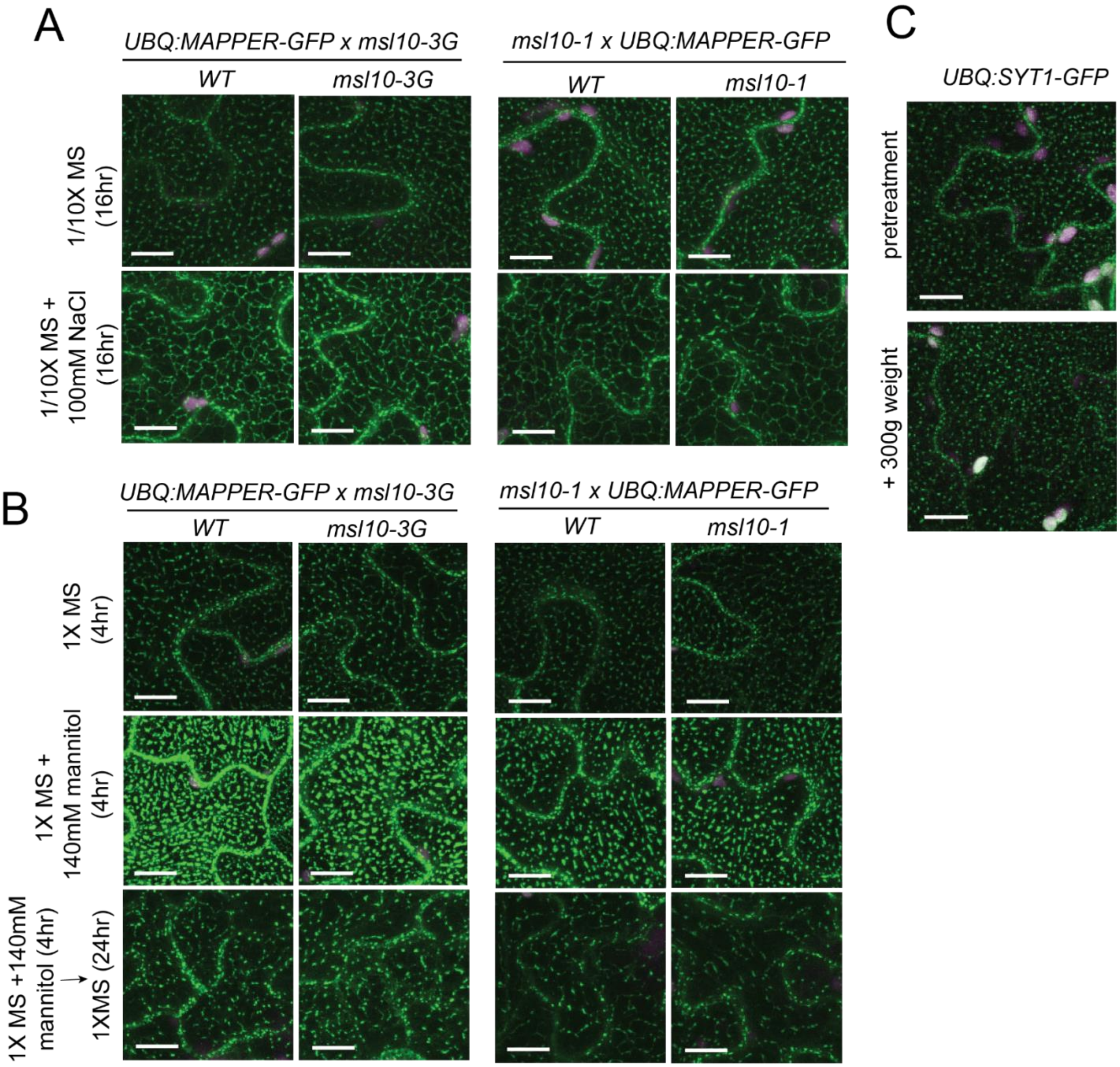


**Figure 3-supplemental figure 1. MSL10 does not influence rearrangements in EPCS morphology in response to osmotic stress in seedlings.** Confocal maximum intensity Z-projections of cotyledon epidermal cells. Green, GFP, magenta, chlorophyll autofluorescence. Scale =10 µm. The seedlings being compared in **(A-B)** are F3 cousins. **(A)** 5-day-old seedlings were transferred from plates and incubated for 16 hr in liquid 1/10X MS or 1/10X MS + 100 mM NaCl. **(B)** Seedlings were grown on 1X MS or 1X MS +140mM plates for 5 days. Seedlings were transferred to liquid media of the same concentration supplemented with 600 nM isoxaben and allowed to equilibrate for 4hr before imaging. MAPPER-GFP puncta size is larger in the presence of mannitol, in contrast to observations in (Lee et al., 2019), and the difference might be attributable to the presence of mannitol in the plates for the entire life of the seedlings, used here. A subset of seedlings incubating in 1X MX + 140 mM mannitol + 600 nM isoxaben were transferred to liquid 1X MS + 600 nM isoxaben to trigger cell swelling, and these were imaged 24hr later. **(C)** 5-day-old seedlings stably expressing UBQ:SYT1-GFP were mounted in water. Cotyledons were imaged before and after a 300 g weight was applied to a 22 x 22 mm coverslip for 20 sec.

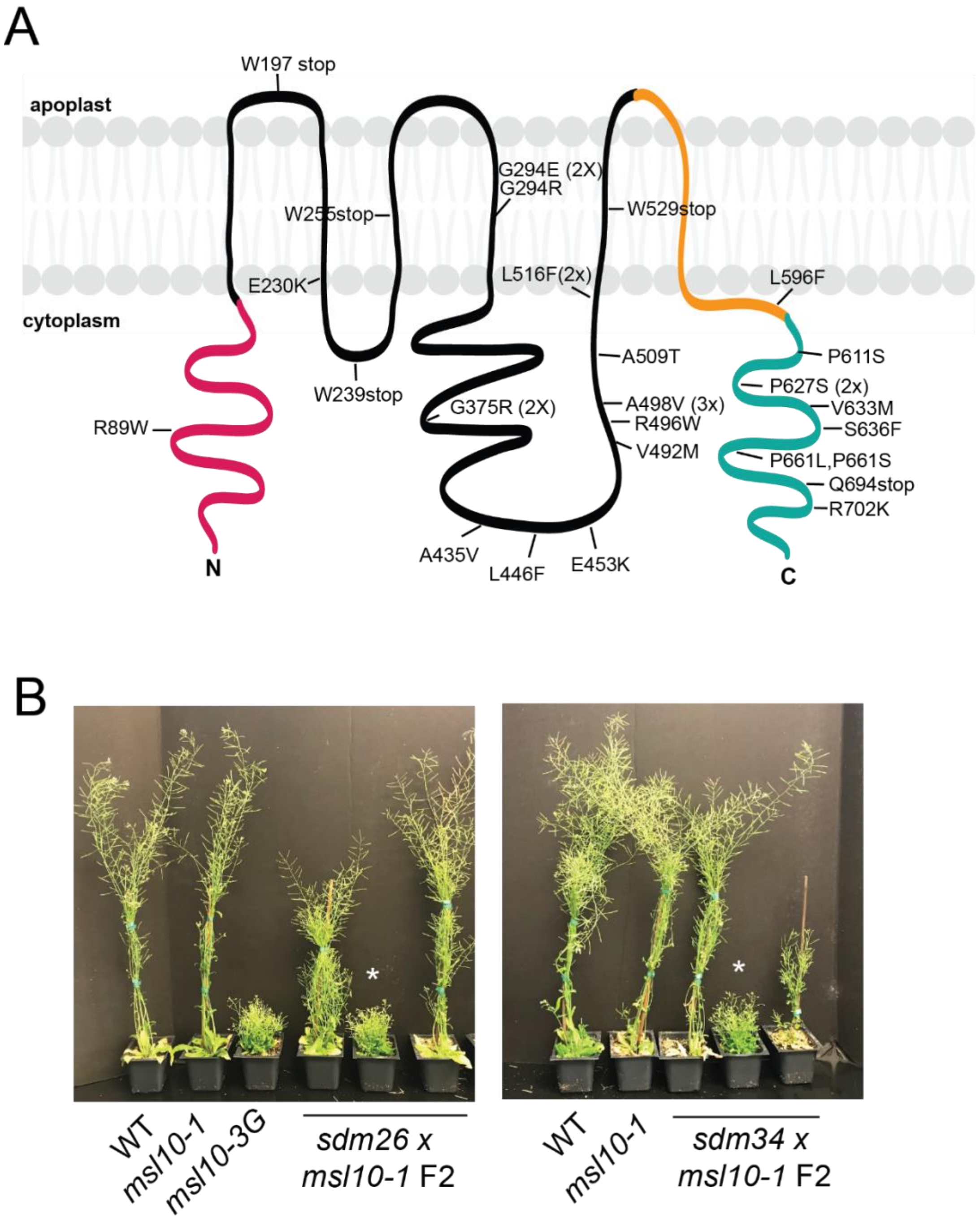


**Figure 4-supplemental figure 1. Intragenic *sdm* mutants and tests to confirm that *sdm26* and *sdm34* causal mutations are extragenic. (A)** Intragenic *sdm* mutations mapped onto the predicted MSL10 topology. In pink is the cytosolic N-terminal domain, in teal is the cytoplasmic C-terminal domain, and in orange is the pore-lining MscS domain. **(B)** Pictures of 5-week old F2 plants. From the *sdm26 x msl10-1* cross, 1 out of 21 F2 plants screened had a dwarf *msl10-3G* phenotype, indicated with the asterisk. From the *sdm34 x msl10-1* cross, there were 2 out of 23 F2 plants that had the dwarfed phenotype. Other F2 plants from both crosses had either a WT or intermediate height. That the *msl10-3G* (dwarf) phenotype could be recovered after crossing to the null *msl10-1* line indicated that the *sdm26* and *sdm34* alleles were not linked to *MSL10*, confirming that they were extragenic suppressors.

A

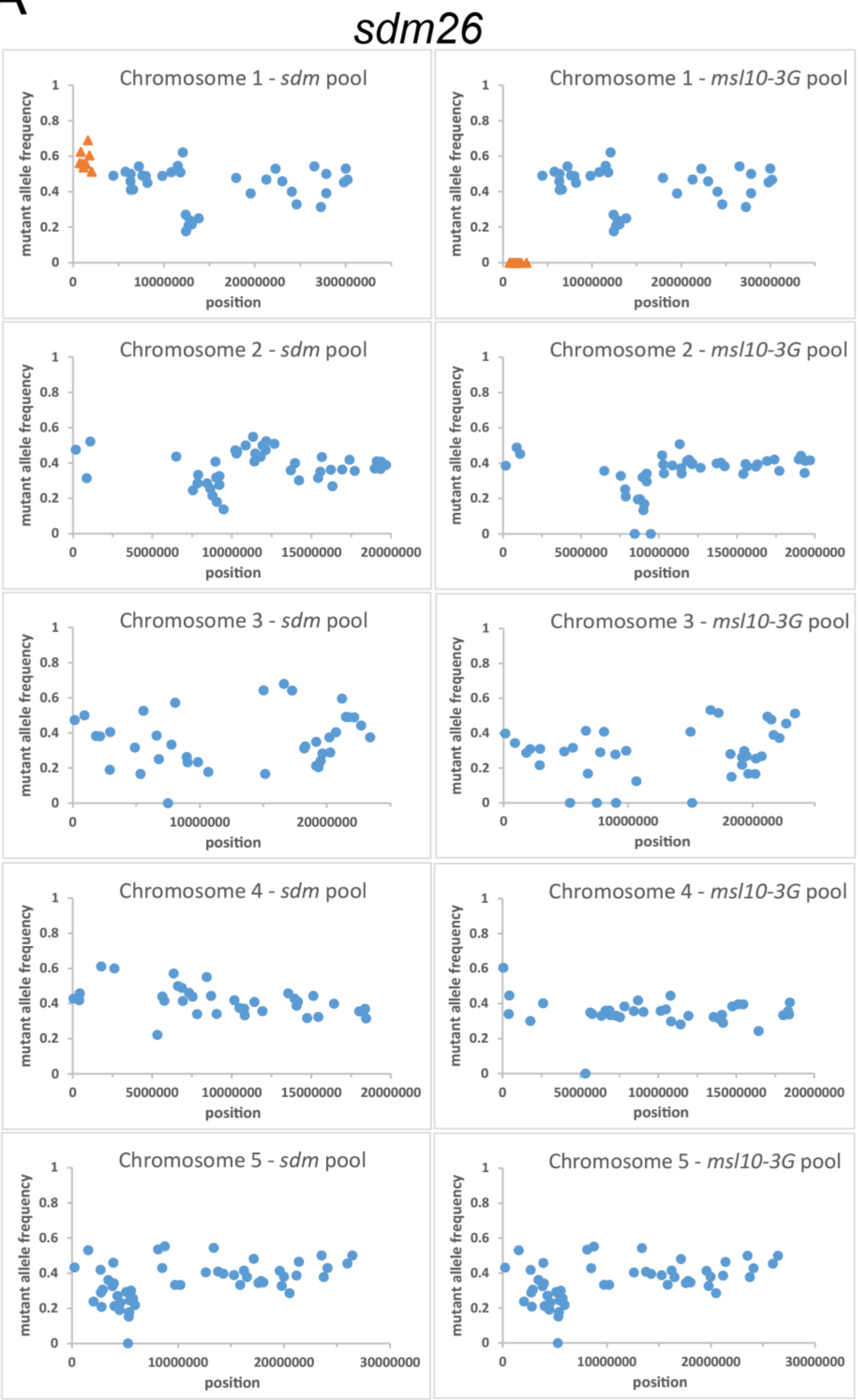


**Figure 5-supplemental figure 1. Mapping-by-sequencing revealed the chromosomal regions containing the causal mutations of *sdm26* and *sdm34*.** *sdm26* and *sdm34* were backcrossed to *msl10-3G*, and the segregating $BC_1F2$ population pooled by phenotype and sent for WGS. **(A,C)** For each SNP identified, the frequency at which this mutant nucleotide was detected compared to the reference nucleotide was calculated and plotted against its chromosomal position. Regions where SNPs are represented with orange triangles were predicted to contain the causal mutation as mutant alleles were absent in the *msl10-3G* phenotypic pool (dwarfed) and present in the *sdm* phenotypic pool (suppressed dwarfing) near the expected frequency of 0.66. **(B,D)** Details of the SNPs in the chromosomal intervals identified in (**A** and **C**). Gene names and functional descriptions were obtained from TAIR.

B

| chromosome | position | reference nucleotide | alternate nucleotide | substitution | locus | gene (description) |
|---|---|---|---|---|---|---|
| 1 | 734512 | C | T | W890stop | AT1G03080 | *NETWORKED 1D* (kinase interacting (KIP1-like) family protein) |
| 1 | 838607 | C | T | P511S | AT1G03380 | *HOMOLOG OF YEAST AUTOPHAGY 18* (yeast autophagy 18 G-like protein) |
| 1 | 1080264 | C | T | A20T | AT1G04140 | unnamed protein (transducin/WD-40 repeat family protein) |
| 1 | 1143747 | C | T | G501S | AT1G04280 | *NAD KINASE-CAM DEPENDENT* (mitochondrial CaM/calcium dependent NAD+ kinase) |
| 1 | 1375564 | C | T | G12E | AT1G04870 | *PROTEIN ARGININE METHYLTRANSFERASE 10* (type I protein arginine methyltransferase) |
| 1 | 1389472 | C | T | D183N | AT1G04910 | unnamed protein (O-flycosyltransferase family protein) |
| **1** | **1625376** | **C** | **T** | **S66F** | **AT1G05500** | ***SYNAPTOTAGMIN 5*** **(endomembrane-localized synaptotagmin)** |
| 1 | 1797131 | C | T | Q30stop | AT1G05920 | unnamed protein (B3 domain protein (DUF313)) |

C

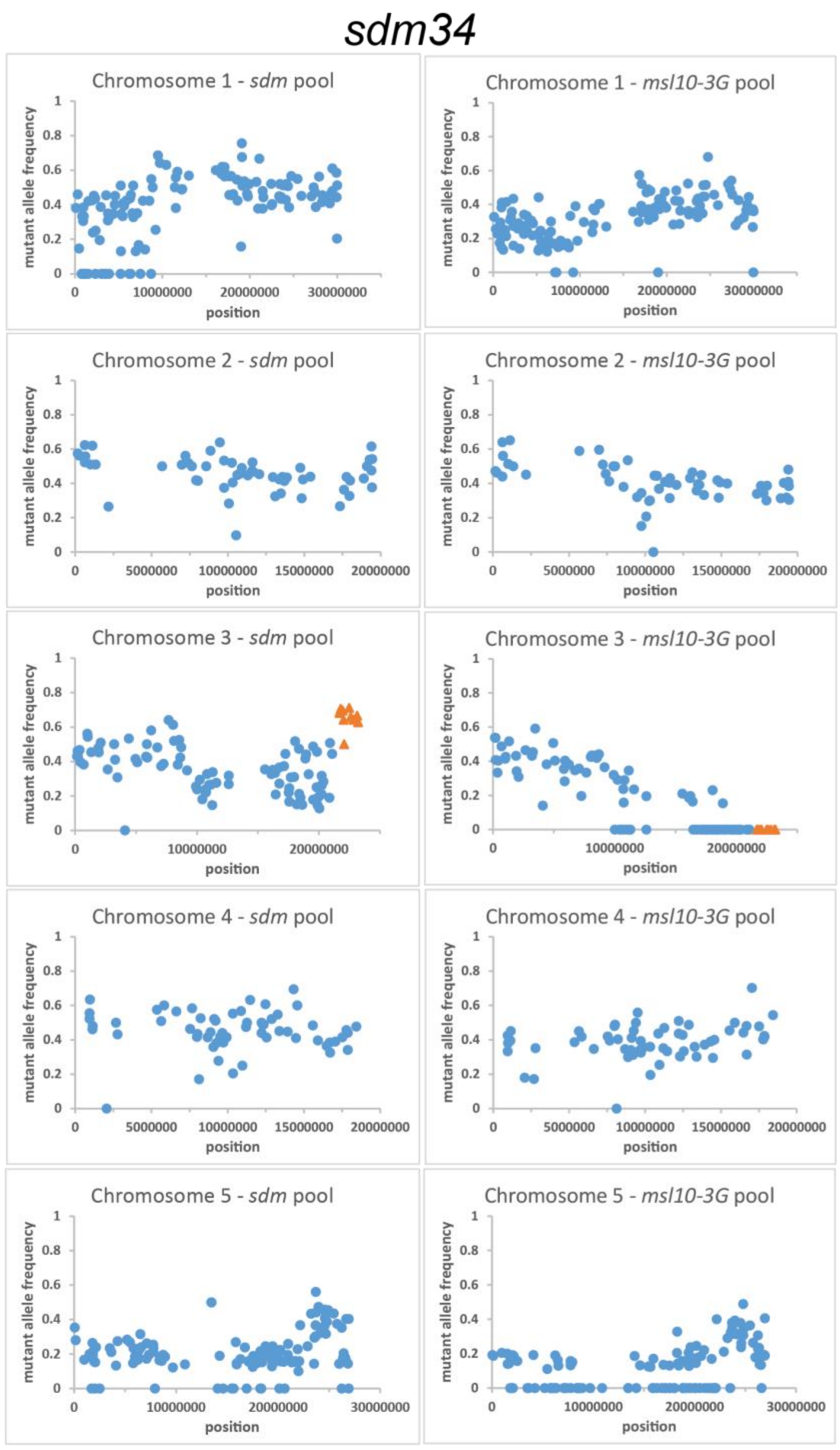


D

| chromosome | position | reference nucleotide | alternate nucleotide | substitution | locus | gene (description) |
|---|---|---|---|---|---|---|
| 3 | 20894222 | G | A | T220I | AT3G56350 | unnamed protein (iron/manganese superoxide dismutase family protein) |
| 3 | 21069109 | G | A | G403R | AT3G56900 | *ALADIN* (component of the nuclear pore complex) |
| 3 | 21672876 | G | A | A242T | AT3G58610 | unnamed protein (ketol-acid reductoisomerase) |
| 3 | 21749070 | G | A | G170E | AT3G58800 | unnamed protein (secretion-regulating guanine nucleotide exhange factor) |
| 3 | 21781019 | G | A | P614S | AT3G58940 | unnamed protein (F-box/RNI-like superfamily protein) |
| 3 | 21956932 | G | A | A915T | AT3G59410 | *GENERAL CONTROL NON-DEREPRESSIBLE 2* (eIF2α kinase) |
| 3 | 22010764 | G | A | D339N | AT3G59580 | *NIN-LIKE PROTEIN 9* (plant regulator RWP-RK family protein) |
| 3 | 22056529 | G | A | A114V | AT3G59710 | unnamed protein (NAD(P)-binding Rossman-fold superfamily protein) |
| 3 | 22472771 | G | A | P109S | AT3G60820 | *PROTEASOME BETA SUBUNIT 1* (encodes 20S proteasome beta subunit) |
| **3** | **22600678** | **G** | **A** | **G427R** | **AT3G61050** | ***SYNAPTOTAGMIN 7*** **(calcium-dependent lipid-binding protein with a C2 domain)** |
| 3 | 22830035 | G | A | S341N | AT3G61690 | *NUCLEOTIDYL TRANSFERASE 8* (putative TNAase) |
| 3 | 23120915 | G | A | E165K | AT3G62500 | unnamed protein (F-box protein FMF) |
| 3 | 23199221 | G | A | P328S | AT3G62710 | unnamed protein (glycosyl hydrolase family protein) |

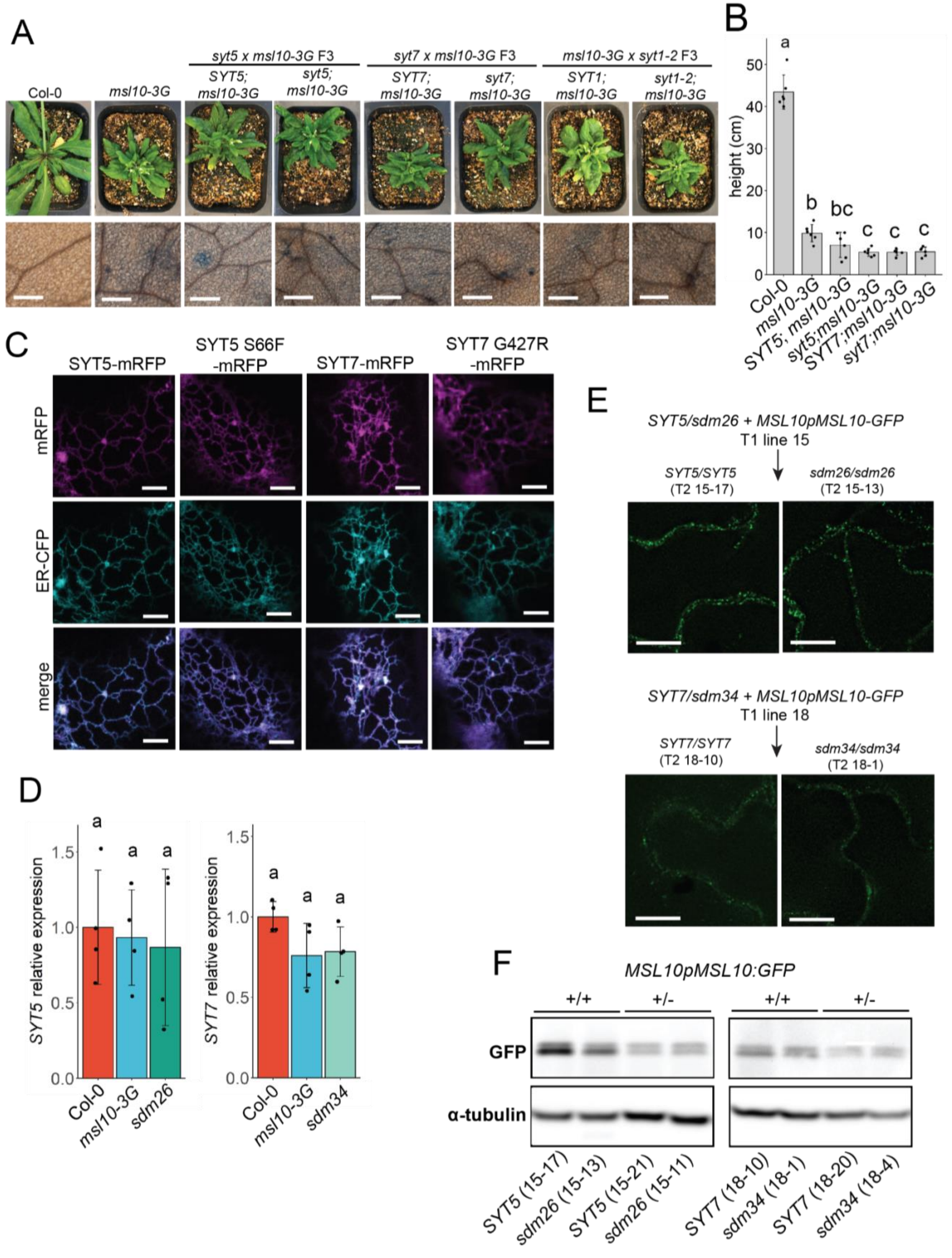


**Figure 6- supplemental figure 1. Null *syt1*, syt5, and *syt7* alleles do not suppress *msl10-3G* phenotypes, and *sdm26* and *sdm34* mutations do not alter SYT5 or SYT7 localization, transcript levels, or MSL10 protein levels.** (A) Plants with the null *syt5*, *syt7*, or *syt1-2* alleles were crossed to *msl10-3G*

plants. Top row: Shown are 4-week-old F3 cousins that are homozygous for the *msl10-3G* allele and homozygous for either the *WT* or null alleles. Bottom row: Trypan blue staining of 4-week-old leaves from the same plants. Scale= 300 μm. **(B)** Height of 6-week-old plants homozygous for both the *msl10-3G* allele and either WT or null *SYT5* or *SYT7* alleles. **(C)** Confocal micrographs of tobacco abaxial epidermal cell transiently co-expressing mRFP-tagged SYT5 and SYT7 constructs under the control of the *UBQ10* promoter and an ER marker (ER-CFP). Scale=5 μm. **(D)** qPCR showing relative *SYT5* and *SYT7* transcript levels *sdm26* and *sdm34* mutants, normalized to *EF1α* abundance using the $2^{-\Delta\Delta Ct}$ method. RNA was extracted from 4-week-old rosette leaves of backcrossed *sdm26* and *sdm34* mutants ($BC_1$F3 cousins). Error bars=SD. Groups indicated with the same letters are not significantly different as assessed by ANOVA with Tukey's post hoc test. **(E-F)** *MSL10pMSL10-GFP* transgenes were introduced into plants heterozygous for the *sdm26 (SYT5 S66F)* or *sdm34 (SYT7 G427R)* alleles (the *msl10-3G* allele had previously been crossed away). Heterozygous T1 plants were identified, and MSL10-GFP stability was compared in T2 siblings that were homozygous for either *SYT* allele. **(E)** Deconvolved image of MSL10-GFP signal in leaf epidermal cells of 4-week-old T2 plants. Scale = 10 μm. (D) Immunoblot of MSL10-GFP protein extracted from 4-week-old T2 siblings. Blots were re-probed with anti-α-tubulin as a loading control.

### Table S1. Primers used in this study.

| primer # | primer name | primer sequence (5' → 3') | purpose |
|---|---|---|---|
| 2229 | LBb1.3 | ATTTTGCCGATTTCGGAAC | genotyping SALK T-DNA insertion lines |
| 3623 | msl10 salk F | GTTGGTTTCTGGGTTTAAGCC | *msl10-1* genotyping |
| 3624 | msl10 salk R | TACTTGGAGTAACCGGTGCTG | *msl10-1* genotyping |
| 702 | MSL10 exon2 For | GCAACGACTAAGGTTTTGCTG | *msl10-3G* genotyping (for CAPS with Taq1 digestion) |
| 663 | MSL10 exon4 Rev | GTTCTTCTTTGTGAGATTAATGTCTTGAGG | *msl10-3G* genotyping (for CAPS with Taq1 digestion), sequencing of *MSL10* genomic DNA |
| 1214 | LB1.SAIL | GCTTTTCAGAAATGGATAAATAGCCTTGCTTCC | genotyping SAIL T-DNA insertion lines |
| 4127 | syt1 genotyping F | GAATTGTCCATGTGAAAGTTGTG | *syt1* genotyping |
| 4128 | syt5 genotyping F | CTGTCAGCGTTTCTCTTAGAG | *syt5* genotyping |
| 4129 | syt5 genotyping R | GAAGAACGTCAACAGTTCAA | *syt5* genotyping |
| 4130 | syt7 genotyping F | GAGAAAGCACTAGATAGTTTGACG | *syt7* genotyping |
| 4131 | syt7 genotyping R | CTGCTGTTTTGCACCATC | *syt7* genotyping |
| 4055 | VAP27-1 For | CACCATGAGTAACATCGATCTGATTG | Amplification of *VAP27-1* ORF for pENTR/D-TOPO cloning |
| 3993 | VAP27-1 Rev | TGTCCTCTTCATAATGTATCCC | Amplification of *VAP27-1* ORF for pENTR/D-TOPO cloning |
| 3988 | VAP27-3 For | CACCATGAGTAACGAGCTTCTCAC | Amplification of *VAP27-3* ORF for pENTR/D-TOPO cloning |
| 4053 | VAP27-3 Rev | TTATGTCCTCTTCATAATGTATCC | Amplification of *VAP27-3* ORF for pENTR/D-TOPO cloning |
| 3990 | SYT1 For | CACCATGGGCTTTTTCAGTACGATAC | Amplification of *SYT1* ORF for pENTR/D-TOPO cloning |
| 3991 | SYT1 Rev | AGAGGCAGTTCGCCACTC | Amplification of *SYT1* ORF for pENTR/D-TOPO cloning / *syt1* genotyping |
| 4038 | ACT8 For | CACCATGGCCGATGCTGATGAC | Amplification of *ACT8* ORF for pENTR/D-TOPO cloning |
| 4039 | ACT8 Rev | TTAGAAGCATTTTCTGTGGACAATGA | Amplification of *ACT8* ORF for pENTR/D-TOPO cloning |
| 4024 | DL1 For | CACCATGGAAAATCTGATCTCTCTGGT | Amplification of *DL1* ORF for pENTR/D-TOPO cloning |
| 4025 | DL1 Rev | CTTGGACCAAGCAACAGC | Amplification of *DL1* ORF for pENTR/D-TOPO cloning |
| 4026 | RAB1c For | CACCATGAATCCTGAATATGACTATTTGTT | Amplification of *RAB1c* ORF for pENTR/D-TOPO cloning |
| 4027 | RAB1c Rev | TTAAGAGGAGCAGCAGCC | Amplification of *RAB1c* ORF for pENTR/D-TOPO cloning |
| 4020 | aCOP1 For | CACCATGTTGACAAAGTTCGAAACC | Amplification of *COPA1* ORF for pENTR/D-TOPO cloning |
| 4052 | aCOP1 Rev | CCGGACTTGAGATGGAGAGCATA | Amplification of *COPA1* ORF for pENTR/D-TOPO cloning |
| 4030 | LOS1 For | CACCATGGTGAAGTTTACAGCTG | Amplification of *LOS1* ORF for pENTR/D-TOPO cloning |
| 4031 | LOS1 Rev | TTAAAGCTTGTCTTCGAAC | Amplification of *LOS1* ORF for pENTR/D-TOPO cloning |
| 4036 | MTO3 For | CACCATGGAATCTTTTTTGTTCAC | Amplification of *MTO3* ORF for pENTR/D-TOPO cloning |
| 4037 | MTO3 Rev | AGCTTGGACCTTGTTAGAC | Amplification of *MTO3* ORF for pENTR/D-TOPO cloning |
| 3986 | AT3G44330 For | CACCATGGCGGAAGAGAAGAAAT | Amplification of *M28 peptidase* ORF for pENTR/D-TOPO cloning |
| 3987 | AT3G44330 Rev | TCCCATTTTCACTTTCCG | Amplification of *M28 peptidase* ORF for pENTR/D-TOPO cloning |
| 4032 | RPT1a For | CACCATGGTGAGAGATATTGAAGAT | Amplification of *RPT1a* ORF for pENTR/D-TOPO cloning |
| 4033 | RPT1a Rev | ATTGTAGACCATATACTTGGG | Amplification of *RPT1a* ORF for pENTR/D-TOPO cloning |
| 4028 | CAT2 For | CACCATGGATCCTTACAAGTATCGTC | Amplification of *CAT2* ORF for pENTR/D-TOPO cloning |
| 4029 | CAT2 Rev | TTAGATGCTTGGTCTCACG | Amplification of *CAT2* ORF for pENTR/D-TOPO cloning |
| 3994 | AT3G62360 For | CACCATGGCGGCCAGTAGGAAG | Amplification of *AT3G44330* ORF for pENTR/D-TOPO cloning |
| 3995 | AT3G62360 Rev | GAACGTCTTCTTTCTAGCAACAGC | Amplification of *AT3G44330* ORF for pENTR/D-TOPO cloning |
| 4022 | RAN1 For | CACCATGGCTCTACCTAACCAG | Amplification of *RAN1* ORF for pENTR/D-TOPO cloning |
| 4023 | RAN1 Rev | CTCAAAGATATCATCATCGTC | Amplification of *RAN1* ORF for pENTR/D-TOPO cloning |
| 3781 | MSL10g upstream seq For | CCCACAGTGTTCTTCTATAATC | Amplification of *MSL10* genomic DNA |
| 3782 | MSL10g downstream seq Rev | CAGTATCACAACGTTTGGTA | Amplification of *MSL10* genomic DNA |
| 699 | MSL10 exon1 For | CAGCACCGGTTACTCCAAGT | Sequencing of *MSL10* genomic DNA |
| 701 | MSL10 exon1 For2 | ACACATTGGACGAAACAGCA | Sequencing of *MSL10* genomic DNA |
| 1611 | MSL10 exon1 Rev | GTTATTGACGTTGAAATTCGCTGCAAGG | Sequencing of *MSL10* genomic DNA |
| 2227 | MSL10 exon3 Rev | CGGACTTCTGAAGTAAGCGCTTATCGGTTTCGTGG | Sequencing of *MSL10* genomic DNA |
| 3789 | MSL10 intron2 Rev | CCATAATTTATCTTTAAAGAATAAAAGCATG | Sequencing of *MSL10* genomic DNA |
| 4145 | SYT5 S66F For | CCTGGGTTGTCTTCTTCGAGCGTCAGAAGTTG | Introducting S66F mutation into *SYT5* by site directed mutagenesis |
| 4146 | SYT5 S66F Rev | CAACTTCTGACGCTCGAAGAAGACAACCCAGG | Introducting S66F mutation into *SYT5* by site directed mutagenesis |
| 4147 | SYT7 G427R For | CAATGGATGCAGTCAGGATGGTGGGAAGTGG | Introducting G427R mutation into *SYT7* by site directed mutagenesis |
| 4148 | SYT7 G427R Rev | CCACTTCCCACCATCCTGACTGCATCCATTG | Introducting G427R mutation into *SYT7* by site directed mutagenesis |
| 4155 | SYT5 For | CACCATGGGTTTCATAGTCGGC | Amplifying *SYT5* for S66F CAPs genotyping / *SYT5* Gateway cloning |
| 4156 | SYT5 internal rev | ACATAAGGCCAGATCTTTGTC | Amplifying *SYT5* for S66F CAPs genotyping / *SYT5* Gateway cloning |
| 4231 | SYT7 dCAPs For | GTAGCACAATGGATGCACTC | Amplifying *SYT7* for G427R dCAPs genotyping |
| 4232 | SYT7 internal Rev | ATCCACTACCGACCGCTC | Amplifying *SYT7* for G427R dCAPs genotyping |
| 4157 | SYT5 Rev | GGAATCACGATAAATTGATTGA | Amplification of *SYT5* for pENTR/D-TOPO cloning |
| 4158 | SYT7 For | CACCATGGGTTTGATTTCTGGG | Amplification of *SYT7* for pENTR/D-TOPO cloning |
| 4159 | SYT7 Rev | CTGCTGTTTTGCACCATC | Amplification of *SYT7* for pENTR/D-TOPO cloning |
| 1758 | attB1-F | ACAAGTTTGTACAAAAAAGCAGGCTCTCCAACCACCATG | Amplifying genes for split-ubiquitin cloning in yeast |
| 1759 | attB2-R | TCCGCCACCACCAACCACTTTGTACAAGAAAGCTGGGTA | Amplifying genes for split-ubiquitin cloning in yeast |
